# Model integration of circadian and sleep-wake driven contributions to rhythmic gene expression reveals novel regulatory principles

**DOI:** 10.1101/2023.08.10.552614

**Authors:** Maxime Jan, Sonia Jimenez, Charlotte N. Hor, Derk-Jan Dijk, Anne C. Skeldon, Paul Franken

## Abstract

Transcriptome studies aim at gaining insight into the molecular pathways underlying biological processes. Analyses of gene-expression dynamics in research on circadian rhythms and sleep homeostasis describe these two processes independently, using separate models such as sinusoidal oscillations and exponential saturating functions. Rhythmically expressed genes are, however, influenced by both processes. We therefore implemented a driven, damped harmonic oscillator model which can accommodate both types of dynamics by varying the degree of damping. This makes it possible to estimate the contribution of circadian and sleep-wake driven influences on the expression of a gene within the framework of a single model. We applied the model to cortex, liver, and blood data obtained in mice and humans. The model reliably captured a wide range of rhythmic dynamics under various experimental conditions, including the long-term amplitude reduction of cortical clock-gene rhythms observed after sleep deprivation. Cortical gene expression was generally influenced more by sleep-wake driven than circadian factors, while the opposite was observed in liver and blood. Importantly, the model suggested that sleep-wake state can alter gene expression with a delayed, long-lasting response not previously considered. Our model further predicted that, perhaps paradoxically, the gain in sleep time after sleep deprivation, delayed re-establishing baseline expression rhythms of intrinsically oscillatory transcripts indicating that similar to insufficient sleep, also excess sleep can impact rhythmic gene expression. Because of the tissue- and gene-specific responses, sleep deprivation led to a profound intra- and inter-tissue desynchronization which in the cortex lasted well beyond phenotypic sleep-wake recovery. The results demonstrate that analyzing rhythmic gene expression must take the complex interactions between circadian and sleep-wake influences into account. The model is a versatile tool with a low number of free parameters to fit and predict gene expression under a variety of conditions relevant to society.

## Introduction

Throughout the brain and body many transcripts exhibit 24h rhythms in gene expression levels [1-3]. These transcriptome rhythms are thought to emerge from cell-autonomous oscillations generated by clock genes engaged in negative transcriptional/translational feedback loops (TTFLs) [4]. The circadian TTFL results in rhythmic expression not only of the clock genes themselves but also that of the numerous other genes they target, many of which are transcription factors thereby setting off daily recurring cascades of transcriptional events comprising the rhythmic transcriptome. Within and among tissue(s) phase coherence is maintained by systemic cues produced by the central circadian clock located in the suprachiasmatic nuclei (SCN) of the hypothalamus, which act as an internal zeitgeber entraining brain and body TTFLs [5, 6]. Transcriptome data have contributed to our current detailed understanding of the molecular architecture of the circadian clock and its tissue-specific functions [7, 8].

Transcriptome studies have also been used in sleep research, in particular to uncover genes and gene pathways implicated in the processes underlying or driven by changes in sleep pressure, which increases while awake and decreases when asleep. These studies have primarily focused on the brain of model species, mainly rats and mice, and used sleep deprivation to experimentally increase sleep pressure. The results showed that sleep-wake states have profound effects on the brain transcriptome [9-12]. By selecting for transcripts that were similarly affected by spontaneous and experimentally induced wakefulness, corrected for the increase in corticosterone levels associated with depriving mice of sleep, we arrived at a short-list of 78 brain transcripts that reliably follow the time course of sleep-wake prevalence both during undisturbed baseline conditions and during sleep deprivation [13]. This short-list features many activity-induced immediately early genes (IEGs) and we observed that their sleep-wake driven dynamics follow exponential saturating functions with time constants similar to those describing the dynamics of delta power [14], a widely used EEG-derived measure gauging sleep pressure. Examples of such transcripts are *Arc* and *Homer1a*, which both play a role in homeostatic down-scaling of synapses, a process considered as one of sleep’s major functions [15-18]. Interestingly, the genes that change their transcription with sleep deprivation include a number of clock-genes [12, 19, 20] which, combined with other observations, suggest a considerable molecular crosstalk between circadian and sleep-wake driven processes in the brain [21]. More recently we found that the brain expression of the core clock-genes *Npas2* and *Clock* followed dynamics similar to that of the sleep-wake driven IEGs and that rhythm amplitudes of all but one of the remaining clock genes showed a long-term reduction following a single, short sleep deprivation [14].

Since under undisturbed conditions the sleep-wake distribution is circadian and because sleep-wake behavior drives the expression of numerous transcripts, many of the genes found rhythmic in circadian transcriptome studies, might oscillate as a consequence of the daily changes in the prevalence of sleep-wake states, and not as a direct consequence of the circadian TTFL within a given tissue. This idea was tested by controlling the time-spent-awake prior to the sampling of cortical tissues at different times of the day. We and others found that under these conditions the majority of rhythmically expressed genes (73-81%) no longer oscillate [15, 22]. Similarly, scheduling sleep in anti-phase with the time it normally occurs in a forced desynchrony protocol, flattened the rhythm of the blood transcriptome in humans, including that of several clock genes [23].

From the above it is clear that sleep-wake driven factors contribute substantially to the circadian transcriptome phenotype in brain and body tissues peripheral to the SCN. Determining which genes and gene pathways are rhythmic as a result of changes in sleep-wake behavior or due to circadian systemic cues, is therefore of interest and of importance when, e.g., assessing the factors underlying the long-term health consequences of circadian misalignment that have been attributed mainly to circadian factors [24, 25]. In a first effort to achieve this, we previously categorized cortical transcripts as either sleep-wake driven or circadian driven using the concepts of the two-process model of sleep regulation [14], a model which stipulates that sleep is regulated by a circadian process (Process C) of sinusoidal shape that interacts with a sleep-wake driven process (Process S) modelled after the dynamics of EEG delta power [26]. In that study [14], we analyzed cortical samples taken over the course of 3 days, i.e., under baseline conditions and during and after a 6h sleep deprivation. The results confirmed that most (63%) of the cortical transcripts rhythmic under undisturbed baseline conditions were categorized as sleep-wake driven when considering the entire 3-day time course. It is, however, unlikely that the rhythmic expression of a given gene is influenced only by either one of the two processes and categorizing genes as such is thus likely to be an oversimplification. Moreover, this approach required model selection among a set of models with different number of free parameters, which is not without issues, and only one type of sleep-wake driven dynamic (i.e., ‘Process S’ type) was considered. Finally, the marked long-term consequences of sleep deprivation on expression dynamics we discovered in that study, especially that of most clock genes, could not be captured by any of the models unless circadian amplitude after the sleep deprivation was altered in the model.

Here we implement a driven, damped harmonic oscillator model to estimate the separate contributions of sleep-wake and circadian processes to the rhythmic transcriptome. In this model circadian systemic cues and sleep-wake driven influences are considered simultaneously as driving factors that effectively accelerate or decelerate peripheral oscillations in gene expression. Importantly, by changing the damping ratio, the model can capture both the dynamics of intrinsically oscillating transcripts (i.e., underdamped in the model) and of overdamped transcripts for which the sleep-wake response approximate exponential saturating functions of Process-S. We applied the model to transcriptome data obtained in mouse cortex and liver tissue, and in human blood and successfully captured the wide range of transcription dynamics observed under conditions of sleep deprivation, forced desynchrony, and a constant routine following 7 days of sleep restriction [14, 23, 27]. The mouse data were used to simulate the effects of sleep deprivation and of recovery sleep on gene expression levels, in particular the time it took for RNA levels to return to baseline, and to estimate within and between tissue desynchronization in gene expression after sleep deprivation. The human data were used to predict transcriptome dynamics during an entire forced desynchrony protocol and during sleep restriction conditions and subsequent constant routine. The results give new insights into the complex interaction between circadian and sleep-wake driven influences on gene expression that might also be relevant for other levels of organization of the rhythmic organism.

## Results & Discussion

### Data sets used to disentangle circadian and sleep-wake dependent influences

Under undisturbed, entrained conditions sleep-wake dependent and circadian contributions to rhythmic gene expression are difficult to disentangle as both factors fluctuate in synchrony with stable phase relationships. To quantify their respective contributions, the timing of sleep (and wakefulness) relative to circadian phase needs therefore to be altered experimentally. In the first dataset used for the current analyses, gene expression in cortex and liver were quantified at 18 time points in mice before (‘baseline’ or ‘BSL’), during, and after (‘recovery’ or ‘REC’) a 6h sleep deprivation (SD; **Fig. 1A**). Sleep-wake behavior was recorded continuously in a separate cohort of mice undergoing the same experimental protocol. The SD kept mice awake at a time-of-day animals are normally mostly asleep, i.e., the first half of the light period (ZT0-6; **Fig. 1A**). The sleep-wake data and cortical transcriptomes were taken from our published and publicly available data [12, 14, 28], while we newly acquired liver RNA-seq data taken from the same mice to assess tissue-specificity of gene-expression dynamics. A second dataset, also publicly available, consists of 2 published experiments quantifying the blood transcriptome in humans using micro-arrays [23, 27]. In the first experiment, participants completed a forced-desynchrony (FD) protocol in which a 28h sleep-wake cycle (and associated dim-light dark cycle) was imposed causing the circadian rhythm to ‘free-run’ at its intrinsic, close-to-24h period. Blood was sampled at 4h intervals during a 28h day when sleep was scheduled at the circadian phase it normally occurs during entrained conditions (‘in-phase’) and during a 28h day when sleep occurred in anti-phase with the circadian cycle (‘anti-phase’; **Fig. 1B**). In the second experiment, participants were given sleep opportunities of either 10h (‘control sleep’) or 6h (‘restricted sleep’) during which they obtained 8.5 and 5.7h of sleep, respectively, for 7 consecutive days preceding a constant routine (CR) during which participants were kept awake for ∼40h with blood samples taken every 3h (**Fig. 1C**). During the CR, light conditions, activity, and food intake were strictly controlled. Before the FD and CR experiments, sleep was recorded at habitual bedtime (‘baseline’; 7.5h of sleep), which we used as the sleep-wake distribution under ‘steady-state’ conditions. While the FD and CR experiments affected timing and duration of sleep-wake behavior, circadian phase, assessed by blood melatonin and cortisol rhythms, remained remarkably unperturbed [23, 29]. This is consistent with analyses of clock-gene rhythms in the mouse SCN which indicated that the central circadian pacemaker is not much affected by changes in the sleep-wake distribution [30-33], although SD has been shown to reduce neuronal activity within the SCN [34]. Furthermore, SD does not alter the phase of circadian activity patterns in mice [35] (but see [36]).

**Figure 1:**
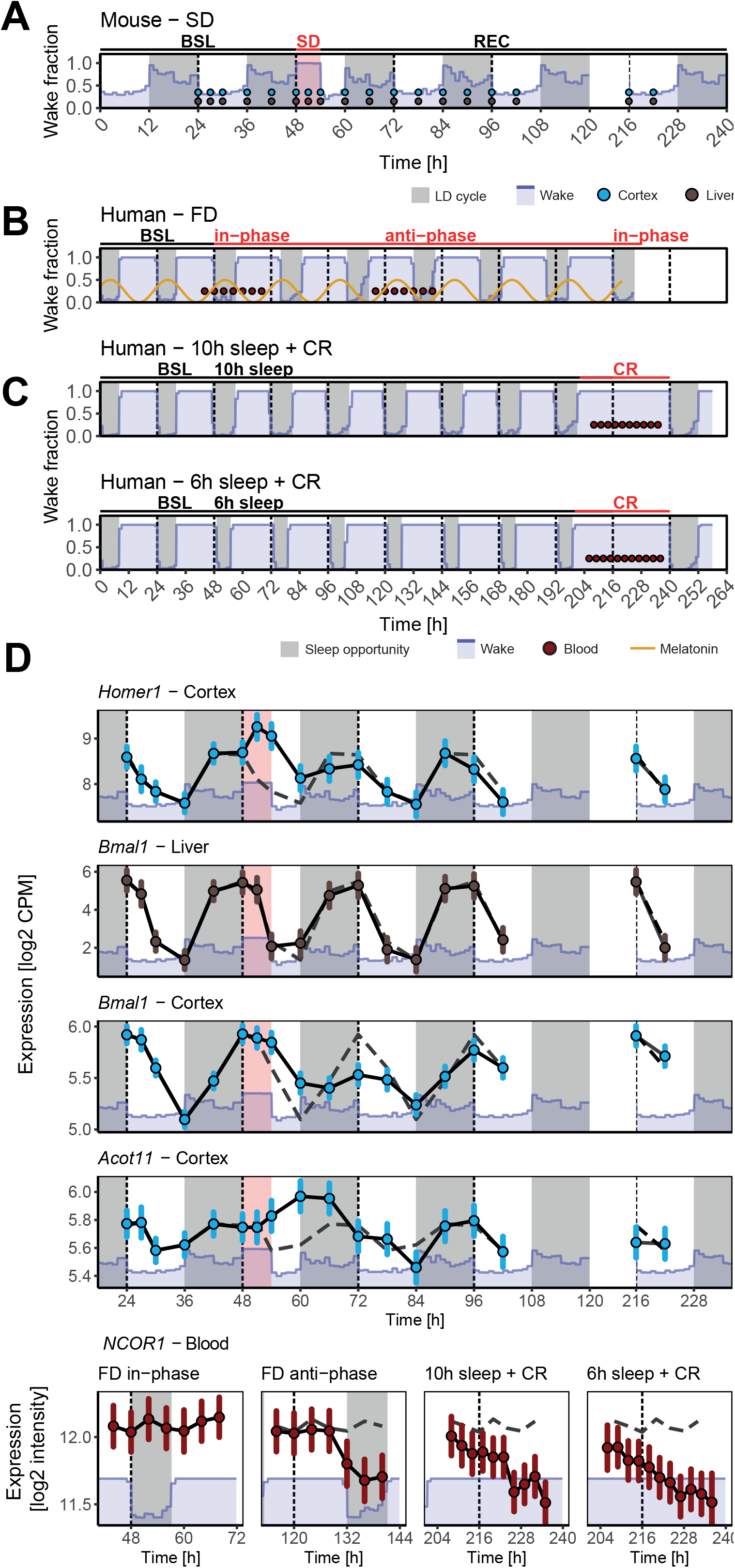
Manipulations of sleep-wake rhythms in mice and humans. **(A)** Sleep deprivation (SD) in mice. Mean fraction of time-spent-awake per hour of recording time (blue line/area, n=12 mice) during baseline (BSL; Days 1 and 2), 6h SD (pink square starting at t=48 on Day 3) and recovery (Days 3-5 and 10). A 2^nd^ batch of mice, undergoing the same experimental protocol, was used for tissue sampling of cortex (blue) and liver (brown points; n=78 mice). Grey background represents the dark periods of the 12h:12h light-dark cycle. Note that the last 2 samples were taken 7 days after the SD. **(B)** Forced Desynchrony (FD) in humans. Mean wake fraction (blue area, n=32) in participants that underwent FD using 28h sleep-wake cycles. Blood samples (red points) were taken during a 28h day when participants slept in-phase and during a 28h day sleep occurred in anti-phase with their circadian melatonin profile. Grey boxes represent scheduled sleep opportunities. **(C)** Constant routine (CR) experiments in humans. Mean wake fraction (blue area, n=36) in participants that underwent a CR after a 7-day control (top panel: ‘10h sleep’, i.e., 8.5h sleep/24h) and a restricted (bottom panel: ‘6h sleep’, i.e., 5.7h sleep/24h) sleep-opportunity schedule. Blood samples (red points) were taken during the CRs. **(D)** Examples of gene expression dynamics in cortex (blue), liver (brown), and blood (red symbols) with mean gene expression (95% confidence interval) per time-point. Solid black lines connect time points, dashed grey lines replicate baseline in mice (before SD) or in-phase dynamics in human. Details as in Panels A-C.

Rhythmic gene expression can follow a dynamic that could be regarded as strictly sleep-wake driven or as strictly circadian driven, illustrated by *Homer1* expression in cortex and *Bmal1* (aka *Arntl*) expression in liver, respectively. *Homer1* expression decreases during the light phase when mice are mostly asleep, increases during the dark when mice are mostly awake, further increases during SD, and quickly (within 18h) re-assumes baseline dynamics during recovery (**Fig. 1D**), with little circadian influence [15]. In contrast, liver *Bmal1* expression oscillates in a regular rhythmic pattern throughout the experiment largely unperturbed by SD (**Fig. 1D**), consistent with *Bmal1* being a core circadian clock gene [37]. Rhythmically expressed genes can, however, show dynamics that do not follow such simple rules [14]. For example, while we find that during baseline the time course of cortical and liver expression of *Bmal1* are similar, SD leads to a substantial and long-lasting reduction in rhythm amplitude during recovery in cortex but not in liver (**Fig. 1D**), demonstrating that, in addition to circadian factors, sleep-wake state affects *Bmal1*’s expression in the former tissue. Furthermore, this amplitude reduction outlasts the effects of SD on recovery sleep [14], indicating that cortical *Bmal1* expression does not seem to simply follow the sleep-wake distribution. Another example is *Acot11*, a gene encoding an enzyme involved in the homeostatic regulation of free fatty-acids [38] and of NREM sleep duration [12]. *Acot11* expression in the cortex increases with SD and also its baseline time course seems consistent with that of a sleep-wake driven gene as it decreases during the light and increases during the dark when animals are predominantly asleep and awake, respectively. Yet, subsequent to SD this relationship appears to invert, as sleep during initial recovery (ZT6-12) is now associated with a strong increase in *Acot11* expression leading to sustained high levels during the subsequent dark phase (**Fig. 1D**). A last example is the dynamics of *NCOR1* expression, which encodes a protein affecting the clock-gene circuitry by acting as co-repressor to the clock-gene *REVERBα* (*NR1D1*) and by activating HDAC3 [39-41]. During the FD, blood *NCOR1* expression appears rhythmic only when sleep occurs in anti-phase with the circadian rhythm (**Fig. 1D**), which might suggest that under normal, in-phase conditions, the sleep-dependent decrease in *NCOR1* expression is opposed by a circadian-dependent increase. However, such a scenario cannot easily explain the important downregulation of *NCOR1* expression with extended wakefulness observed during the two CRs in the second experiment (**Fig. 1D**).

These examples illustrate that rhythmic gene expression results from an often complex interaction between the responses to circadian and sleep-wake dependent drives that seem to greatly differ among genes and tissues. It also illustrates the difficulty to reconcile a gene’s dynamics under different experimental protocols. Quantifying and comparing the relative importance of these factors in driving the rhythmic transcriptome requires a novel modeling approach integrating sleep-wake and circadian dependent influences on gene expression.

### Rhythmic gene expression as a driven, damped harmonic oscillator

Transcriptome rhythms measured in peripheral organs are thought to arise from transcriptional-translations feedback loops (TTFL) made up of the core circadian clock genes [4]. According to this scenario, local tissue rhythms are kept in phase with each other and with the light-dark cycle by signals generated by the SCN which take the role of an internal zeitgeber. At the same time, the SCN drive rhythms in overt behaviors such as sleep and wakefulness [**Fig. 2A – left** [42, 43]]. Although perturbations of sleep are known to impact gene expression, including that of clock genes, only a hand-full of studies have considered the influence of the sleep-wake distribution on the rhythmic transcriptome [9, 14, 23, 27]. Most studies only examine the immediate effect of SD or assess the interaction of sleep-wake and circadian driven processes using experimental protocols such as ‘around-the-clock’ SDs [15, 22]. In such protocols it remains, however, unclear whether residual rhythmicity is caused by circadian factors, including time-of-day differences in the response to SD, or by differences in sleep-wake history prior to the SDs. Similarly, modeling sleep-wake driven dynamics using exponential saturating functions following the example of the dynamics of EEG delta power [14, 44] does not include a circadian component, and interactions between circadian and sleep-wake related factors, beyond simple additive effects, have not been not considered (**Fig 2A - middle**). The model we propose allows for such interaction and provides a framework to quantify the relative contribution of circadian and sleep-wake dependent factors on rhythmically expressed genes. These genes can be modeled as intrinsically rhythmic, i.e., because they are closely associated with the circadian TTFL, or they can appear rhythmic because they follow circadian and/or sleep-wake dependent drives but, in the absence of such recurring drives, do not oscillate (**Fig. 2A - right**). We have used earlier implementations of this modeling approach to simulate the effects of sleep-wake state on *Per2* mRNA and protein levels [45, 46].

**Figure 2:**
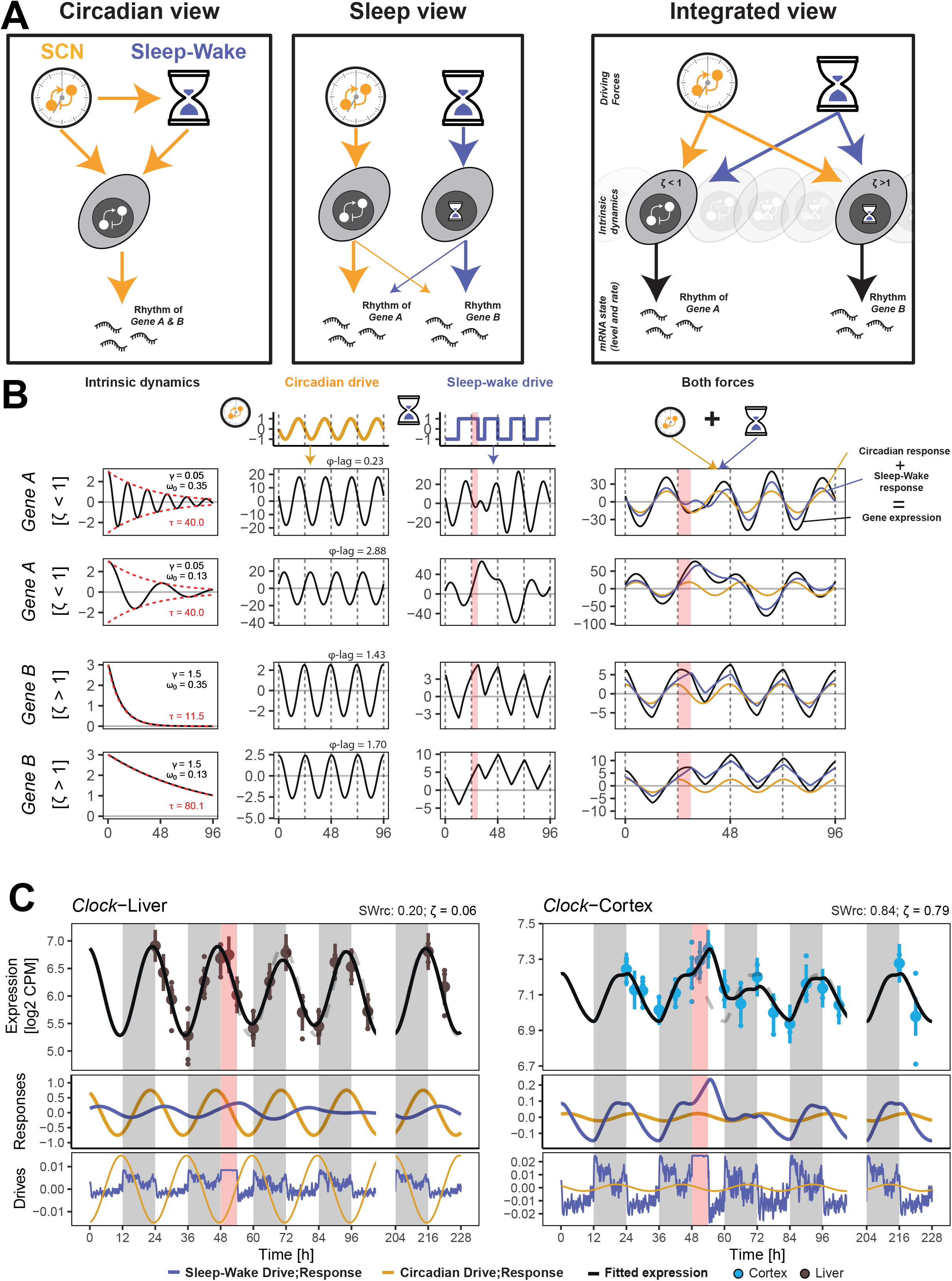
Modeling gene expression using a damped driven harmonic oscillator. **(A)** Schematic of circadian view of generation of rhythmic gene expression (left) in which the SCN directly or indirectly drives or entrains oscillations of gene expression generated by local circadian clocks (TTFL) in peripheral cells. Sleep view (middle) separates circadian and sleep-wake related genes, each regulated by different dynamics. The integrated view (right panel) considers each gene to be regulated to a varying degree by systemic circadian and/or sleep-wake dependent influences which act as drives on gene expression in the periphery. **(B)** Illustration of the damped driven harmonic oscillator model. According to a gene’s intrinsic properties, two types of expression dynamics can be observed when expression is removed from equilibrium and no drive is applied: an underdamped system oscillating with a decaying amplitude (upper panels, hypothetical *Gene A*, damping ratio ζ < 1) and an overdamped system (bottom panels, *Gene B*, ζ > 1) where expression returns to equilibrium position without oscillation according to exponential decaying function (red-dashed lines) with a time constant τ determining the time it takes to recover. τ depends on ζ and the natural frequency, ω_0_. For each gene, examples of two ω_0_ values are given: 0.35 and 0.13 [rad/h], illustrated in the upper and lower row panels, respectively. External recurring driving factors are required to maintain gene expression entrained and rhythmic (circadian drive in yellow, sleep-wake drive in purple; middle two panels). The difference between ω_0_ and the frequency of the external drive determines the phase-lag (φ-lag) between drivee and response. Combing the responses to each drive generates the observed rhythm in gene expression (right panels). Pink areas represent sleep deprivation. **(C)** Model fit for expression of *Clock* in liver (left) and cortex (right panels). Circadian (yellow) and sleep-wake (purple) drives applied on the model (bottom), circadian and sleep-wake responses to the drives giving the best fit (middle), fitted expression in black with mean gene expression (95% confidence interval, upper panel). Dashed grey lines replicate baseline. *SWrc* is the relative contribution of the sleep-wake response (see Results).

The measured level of the expression of a gene at a given time point reflects the net result of mRNA synthesis and degradation. With our data we cannot assess whether changes in gene expression resulted from changes in production, degradation, or both. In the following we nevertheless use the terms synthesis and degradation when referring to net increase and decrease in mRNA levels, respectively. We propose a simple framework in which we suppose that the level of mRNA of a gene is *X(t)* where *t* is time. We suppose that the rate of synthesis of *X(t)* will depend on intra-tissue factors such as the levels and activity of transcription factors, temperature, and metabolites affecting mRNA regulation, which we group together in a single ‘tissue environment’ variable *Y(t)*. We suppose the rate of degradation depends on the level of *X(t)*. In a simplest (linear) approximation, the rate of change of mRNA may be written as

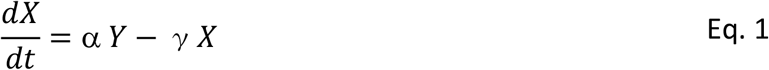

where α describes the effect of the tissue environment on the synthesis rate of *X(t)* and *γ* is the degradation rate per unit *X(t)*. We assume that the tissue environment variable is affected by external factors *F*(*t*) such as the circadian and sleep-wake drives and that there is feedback between the gene of interest and the tissue environment so that

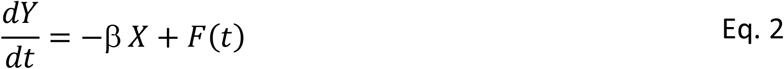

where β describes the strength of the feedback between the gene of interest and the tissue environment.

We let *X*(*t*) = *X*_*b*_ + *x*(*t*), *Y*(*t*) = *Y*_*b*_ + *y*(*t*) and, *F*(*t*) = *F*_*b*_ + *f*(*t*) where *X*_*b*_, *Y*_*b*_, and *F*_*b*_ are fixed baseline values that satisfy equations (Eq. 1) and (Eq. 2) when 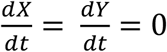. Then substituting for *X(t)* and *Y(t)* in equations (Eq. 1) and (Eq. 2), differentiating (Eq. 1) with respect to time and substituting in for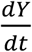 from equation (Eq. 2) leads to the equation for a damped harmonic oscillator (Eq. 3) (see the Supplementary Material for further details).

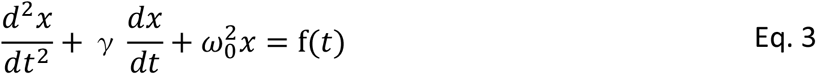

where 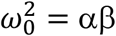 and f(*t*) = α *f*(*t*). In this equation, *x*(*t*), represents the level of mRNA of a gene quantified as normalized counts from RNA-sequencing (in log2 counts per million or CPM) for the mouse tissues or from *Affymetrix* microarrays (in log2 probe intensities) for human blood samples. The term 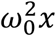 arises from the feedback between the gene and its environment and could be viewed as, e.g., an auto-inhibition through negative feedback [47], as is the case for the expression of clock genes that comprise the circadian TTFL. A large value of 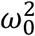 translates into a strong negative feedback controlling gene expression. In contrast, a weak negative feedback will result in gene expression rhythms being driven mostly by changes in external factors. Another intrinsic factor determining gene expression dynamics is the degradation constant, *γ*, which opposes changes in gene expression and introduces a time delay in response to external driving factors.

The model can capture both intrinsically oscillatory and non-oscillatory genes. Using the standard terminology of simple harmonic oscillators in the absence of time dependent external driving factors (f(*t*) = 0), when the damping ratio, ζ = *γ*/2ω_0_ < 1;, the oscillator is said to be underdamped. When released from a position away from equilibrium, the expression of the hypothetical gene, *Gene A*, will oscillate around equilibrium with an amplitude that decreases on a timescale determined by damping constant *γ* (**Fig. 2B - top two rows**). However, when ζ> 1 (i.e., overdamped), gene expression will not oscillate and reverts to the equilibrium directly (hypothetical *Gene B*; **Fig. 2B - bottom two rows**). For underdamped genes, the time required for the expression to return to equilibrium (τ) is determined by *γ*, while for overdamped genes it depends on *γ* and ω_0_ (Eq. 4; **Fig. 2B - red line**).

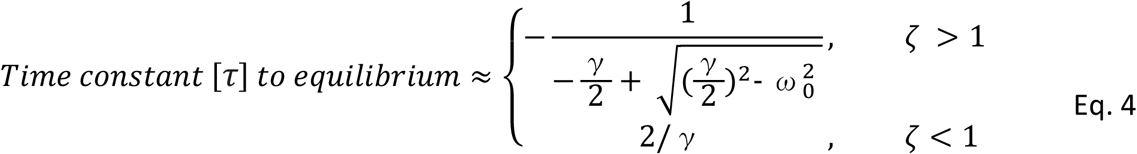

Recurring external driving factors (f(*t*) in Eq. 3) are needed to assure phase coherence of the daily transcriptome changes among and within tissues and, if *γ* > 0, to maintain rhythmicity. Such external factors can either follow continuous oscillations (**Fig. 2B - 2**^**nd**^ **column**) originating, for example, from the SCN or result from discrete physiological or behavioral events such as being (kept) awake or asleep (**Fig. 2B – 3**^**rd**^ **column**), which in this schematic includes a SD (pink bars). We refer to these two types of driving factors as ‘circadian driven factor’ (*f*_*C*_ (*t*)) and ‘sleep-wake driven factor’ (*f*_*SW*_ (*t*)), respectively. In the model we base *f*_*SW*_ on the fraction of sleep (*S*(*t*); i.e., NREM + REM sleep) and wakefulness (*W*(*t*)), measured within a given time interval, *t*, multiplied by their respective coefficients, *β*_*s*_ and *β*_*w*_ (Eq. 5, see Methods). The circadian drive, *f*_*C*_ (*t*), is modeled as a sinewave with a 24h period and a free phase and amplitude (φ and *A*; Eq. 5).

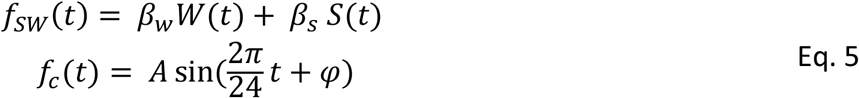

Together these two factors affect the rhythmic expression of a gene by increasing or decreasing its acceleration i.e. the rate of change of its synthesis rate.

The combined effect of the two driving factors on the oscillator can be mathematically decomposed into the responses to either factor (see the Supplementary material). Summing the separate contributions again reconstructs the gene-expression dynamics fitted by the model (**Fig. 2B - right column**). In the **Figure 2B** schematic the relative contributions of the two driving factors (and their respective responses) to the expression dynamics of *Genes A* and *B* are similar in magnitude prior to SD, yet because of their different intrinsic properties, the response to the same sleep-wake perturbation can considerably differ. Besides ζ, the response also depends on the phase-lag between the oscillator and the drive which is determined by the frequency ratio (*r* = ω/ω_0_) between the frequency of the drive 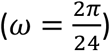 and the natural frequency (ω_0_) (Eq. 3). If *r* = 1, the phase-lag is 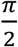 and the oscillator is said to be in resonance. If *r* ≫ 1, the phase-lag increases and an inertia in the response of the oscillator is observed such that the rate of gene expression will only slowly change after a change in the external drive. In contrast, when *r* ≪ 1, the phase-lag decreases, causing the rate of gene expression to change already before the external driving factors can exert their influence, because of the feedback generated by the system.

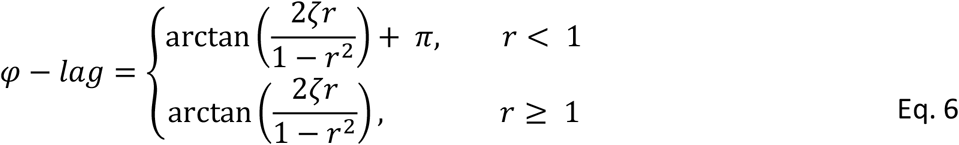

With different contributions from the two external driving factors and different intrinsic parameters, the model can capture a large variety of dynamics (**Fig. 2B - right column**).

The parameters *γ*, ω_0_, *β*_*w*_, *β*_*s*_, *A*, and φ of the model were estimated by fitting gene expression in mouse cortex, liver and human blood (see Methods). The parameters were estimated independently for each gene and tissue (see **Supplementary Table 1)**. While the model fitted gene expression at the time points the tissues were sampled, with the optimized parameters the model was then used to predict the entire time-course when sleep was recorded, including for example the habitual bedtime (BSL) recording prior to the FD protocol as well as all days during that protocol.

**Figure 2C** illustrates the responses to the two driving factors the model estimated for the expression dynamics of *Clock* with strikingly different results in the two tissues. As for *Bmal1* (**Fig. 1D**), *Clock* expression in the liver displays a sinewave oscillation unperturbed by SD. In contrast, cortical *Clock* expression decreased when animals were asleep, increased when awake spontaneously and during SD (**Fig. 2C**). Although the model fitted the *Clock* expression dynamics equally well in the two tissues (Kendall’s τ = 0.56 and 0.73 in cortex and liver, respectively) the damping ratio greatly differed (ζ = 0.79 and 0.06, respectively). Of note, we used Kendall’s τ as an estimate of goodness of fit for time series [48] because R^2^ is inadequate for nonlinear regression [49]. In liver, *f*_*C*_ and its response was much stronger than that of *f*_*SW*_ while the opposite was observed in the cortex where *Clock* dynamics resembled that of a sleep-wake driven gene such as *Homer1* (**Fig. 1B**) [14]. We quantified the relative contribution of the two drives by calculating a *SW-response contribution* (*SWrc*) metric as follows: the peak-to-trough amplitude of the response to *f*_*SW*_ (*A*_*SWr*_) in baseline was expressed as a fraction of the peak-to-trough amplitude of the summed response to the 2 forces (*A*_*SWr*_ + *A*_*Cr*_; Eq. 7). *SWrc* can vary between 0 and 1 with 0 indicating that the summed response is entirely due to *f*_*C*_, 1 to *f*_*SW*_, and 0.5 indicating equal contributions.

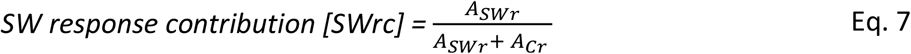

We determined this fraction under undisturbed baseline conditions because *SWrc* depends on the sleep-wake distribution and thus will be larger during, e.g., SD. For the expression of *Clock, SWrc* in liver was 0.20 and in cortex 0.84 (**Fig. 2C**), reflecting well the circadian and sleep-wake driven nature of the dynamics in the two respective tissues, comparable to *SWrc* values obtained for *Bmal1* expression in liver (0.10) and *Homer1* in cortex (0.83; **Fig. 3**).

**Figure 3:**
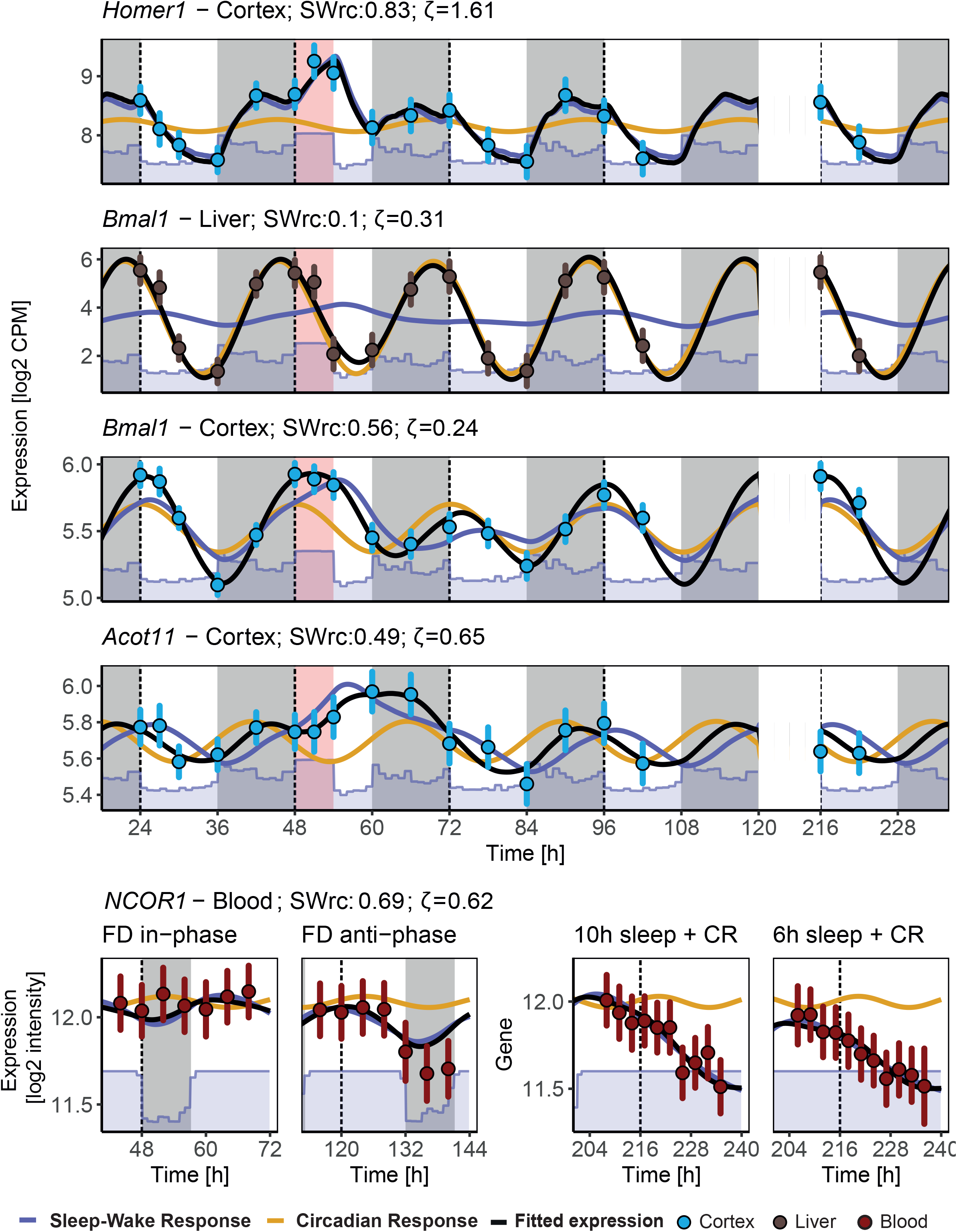
Model fits for the gene-expression examples in Figure 1D. Fitted dynamics (black line) of **c**ortical *Homer1* expression follows almost exclusively the sleep-wake response (purple line) while *Bmal1* in the liver the circadian response (yellow line). *Bmal1* and *Acot11* in cortex and *NCOR1* in blood follow a combination of a sleep-wake and circadian response. SWrc: Sleep-wake relative contribution. Details as in **Fig. 1D**.

It is important to note that i) with the terms circadian and sleep-wake driven we here refer only to the type of drive the expression of a particular gene responds to and not whether the gene can intrinsically display rhythms or not (i.e., is over-or underdamped), ii) the oscillator’s response does not only depend on the sign and magnitude of the exerted drives and the gene’s intrinsic properties (i.e., ω_0_ and *γ*), but also on the state of the oscillator, such as the expression level and the rate at which it changes at the time the drive is applied, and iii) although the model can easily differentiate genes as being over-or underdamped when their expression responds to the sleep-wake distribution, purely circadian driven genes that are under-or overdamped will display indistinguishable dynamics (**Fig 2B - 2**^**nd**^ **column panels in orange**). Assessing this would require experimentally changing the magnitude of the circadian drive.

Our model not only reliably captured straightforward gene expression dynamics but also less predictable scenarios. In the simplest scenario, the rhythmic expression of a ‘pure’ sleep-wake driven gene will tightly follow the sleep-wake distribution, independent of circadian phase (or time-of-day), and the gene will be intrinsically overdamped (non-oscillatory, ζ > 1), together resulting in dynamics approximating those following exponential functions such as observed for many immediate-early genes [IEGs [14]], including *Homer1* (**Fig. 3**), and for EEG delta power (**Fig. S1**, see also **Supplement text**). On the other hand, the expression of a ‘pure’ circadian-driven gene will continue oscillating because it is intrinsically underdamped (oscillatory; but see comment in previous paragraph) and responds only to circadian drives (i.e., with a low *SWrc*) such that amplitude and phase are unaffected by changes in sleep-wake state as was observed for *Bmal1* and *Clock* in liver (**Fig. 3, Fig. 2C**). The model established that for the 3 remaining genes highlighted in **Figure 1** the sleep-wake and circadian drives contributed approximately equally to their expression dynamics (*SWrc*: 0.49-0.69). Yet, because of their different intrinsic properties (**Table 1**), expression dynamics responded very differently to the drives applied. Wakefulness was found to apply a positive drive accelerating cortical *Bmal1* expression. Nevertheless, its expression did not increase during SD because the natural frequency is close to the baseline sleep-wake frequency (ω_0_ = 0.21 and 0.26, respectively) and thus the sleep-wake response is close to *Bmal1*’s maximum amplitude, and because the circadian response decreases during the SD. The model found that the prolonged amplitude reduction of *Bmal1*’s oscillation in the cortex after SD resulted from a combination of a low damping constant (*γ*), which increased the time to return to equilibrium (τ = 20*h*, Eq. 4), and the reduction in time-spent-awake during the recovery dark period, which reduced the normal increase in gene expression rate at this time-of-day. Wakefulness also accelerated the rate of cortical *Acot11* expression (**Fig. 3**). The model found that the peculiar, prolonged increase in *Acot11* expression during recovery sleep was due to a weak negative feedback 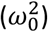 and thus a long phase-lag between drive and response. This inertia to the wake drive during SD was strong as it would have required 2h of continuous sleep to counter it and for sleep-wake response (blue line) to start decreasing. In addition to this inertia, the interaction between the circadian and sleep-wake responses maintained a high expression for 9h after SD, further delaying a reduction of *Acot11* expression. In contrast to the two previous examples, wakefulness decelerated the rate of *NCOR1* expressing in human blood. The model suggested a weak negative feedback to underly the continued decrease in *NCOR1* expression for the entire duration of both CRs. This result highlights that the contribution of the sleep-wake response and the circadian response depend on the experimental condition: in baseline the two contributions were similar (*SWrc* = 0.59) but in anti-phase thereby flattening gene expression while during the CRs, when subjects are kept awake for 40h, the sleep-wake contribution becomes larger relative to the circadian contribution (*SWrc* = 0.90 during CR). These examples also underscore that a gene’s expression can appear rhythmic for a variety of reasons which can greatly differ according to tissue. Moreover, the circadian and sleep-wake driven influences on the expression of some genes can be revealed only during longer-term sleep disruptions and would have gone unnoticed under undisturbed conditions. Our strategy importantly differs (and captures other genes) from simply assessing differential expression immediately after the SD, which has been used to categorize a gene as sleep-wake driven (**Fig. S2**). Finally, with these examples the model revealed that sleep-wake driven responses can importantly deviate from the dynamics following exponential saturating functions that are typically associated with sleep-wake driven responses.

**Table 1:** Parameters estimated for the expression of the genes in Figs. 2C and 3A, with **x** as gene expression (log_2_ of CPM or probe intensity), ***γ*[h**^**-1**^] the daemping coefficient, and ω_0_**[radian * h**^**-1**^] the natural frequency of the oscillator. **β** _**w**_ **[x * h**^**-2**^ **\* W**^**-1**^] and **β** _**S**_ **[x * h**^**-2**^ **\* S**^**-1**^] are the wake and sleep coefficients for *f*_*SW*_ with **W** and **S** as the wake and sleep fraction. **A [x * h**^**-2**^] is the amplitude and **φ [radian]** the phase of *f*_*C*_. Derived parameters of the model are: Damping ratio of the oscillator **[ζ]**, time-constant **[τ;** Eq. 4] to return to equilibrium, **[phase-lag;** Eq. 6] between driving forces and oscillator phase, Sleep-wake response contribution **[SWrc;** Eq. 7].

|  |  | *Homer1* | *Bmal1* |  | *Acot11* | *NCOR1* | *Clock* |  |
| --- | --- | --- | --- | --- | --- | --- | --- | --- |
|  |  | Cortex | Liver | Cortex | Cortex | Blood | Liver | Cortex |
| Optimized parameters | $\gamma$ | 1.68 | 0.12 | 0.10 | 0.18 | 0.10 | 0.03 | 0.42 |
| | $\omega_0$ | 0.52 | 0.19 | 0.21 | 0.14 | 0.07 | 0.22 | 0.27 |
| | $\beta_w$ | 0.450 | 0.020 | 0.015 | 0.013 | -0.003 | 0.008 | 0.024 |
| | $\beta_s$ | -0.510 | -0.030 | -0.017 | -0.015 | 0.006 | -0.006 | -0.031 |
| | $A$ | 0.050 | 0.100 | 0.006 | -0.007 | 0.002 | 0.010 | 0.002 |
| | $\phi$ | 3.42 | 4.59 | 3.78 | 2.50 | 3.18 | 4.85 | 2.51 |
| Derived parameters | $\zeta$ | 1.61 | 0.31 | 0.24 | 0.65 | 0.62 | 0.06 | 0.79 |
| | $\tau$ | 5.47 | 16.73 | 19.26 | 10.82 | 20.80 | 78.82 | 4.68 |
| | $\phi - lag$ | 1.13 | 2.37 | 2.27 | 2.35 | 2.76 | 2.80 | 1.52 |
| | $SWrc$ | 0.83 | 0.10 | 0.56 | 0.49 | 0.69 | 0.20 | 0.84 |

### Assessing the model’s performance against alternative models

Before characterizing the dynamic properties of the full transcriptome, we evaluated the performance of the model and possible overfitting by comparing it to both simpler and more complex models considering all datasets. The evaluation was performed on the subset of genes and probe-sets that showed rhythmic expression during baseline for mice and when sleep occurred in phase with melatonin production for humans. The selection of this rhythmic subset was necessary as our model aims at capturing the dynamics of rhythmic genes and fitting arrhythmic or very noisy genes would automatically favor less complex models. As mentioned earlier, ‘pure’ sleep-wake driven and ‘pure’ circadian-driven genes can display undistinguishable rhythmic patterns in baseline. Both categories of genes can thus be captured in an unbiased fashion with a simple sinewave fit and independently of their response to sleep perturbation. The time courses of the top 1000 most significant ‘sinusoidal’ genes per tissue (cortex, liver, and blood) were used to assess the model’s performance, i.e., a total of 3000 genes.

Our model has 6 free parameters (k=6 [γ, ω_0_, *β*_*s*_, *β*_*w*_, φ, *A*]; see Eq. 3 and Eq. 5), with the equilibrium position (intercept) fixed to the mean gene expression in baseline in mouse and in-phase data in human. The model integrated the two human transcriptome experiments as one and model parameters were simultaneously optimized such that, e.g., 1 minute of wakefulness in the FD protocol has the same accelerating effect as 1 minute of wakefulness during the CRs following the control- and restricted-sleep conditions. We did, however, allow different intercepts between the FD and the CRs after the control- and restricted-sleep conditions (k=7).

To evaluate the fit and complexity of our model (Hypothesis 1 or H_1_) we contrasted it to the following 4 alternative models (H_A_): i) a linear regression model based on independent fixed effects for each time-point (k=18 and 35 in mouse and human, respectively) known to over-fit the data [14], ii) the oscillator model with a sleep-wake drive only or, iii) with a circadian drive only (k=4 and 5), and iv) a simple additive model in which a fixed circadian effect (sinewave) is added to a sleep-wake effect without intrinsic dynamics integrating these effects (k=5 and 6; see Methods). We compared the Bayesian Information Criterion (BIC) statistic of each of the 4 H_A_ models to that of H_1_. The BIC considers the model’s goodness of fit while penalizing for complexity. A ΔBIC was calculated for each of the 4 comparisons with positive values indicating support for H_1_ and negative values indicating support for H_A_. In general, the ΔBIC indicated more genes with a better fit for H_1_ over both simpler and more complex models (ΔBIC>0: 97, 61, 88, and 68% of all 3000 genes, for H_A_ i-iv, respectively), even when using a more stringent ΔBIC (>2: 97, 55, 85, and 63%, respectively; **Fig. 4A - top**). In some cases, ΔBIC favored H_A_, although a strong support was found only for a minority of genes or probes (ΔBIC<-2: 2, 30, 7, and 24%, respectively). It shows that despite having far fewer parameters than the linear model with independent time-effect, goodness of fit for H_1_ is still high (∼0.1 ΔKendall’s τ) and is improved compared to simpler models (**Fig. 4A - bottom**).

**Figure 4:**
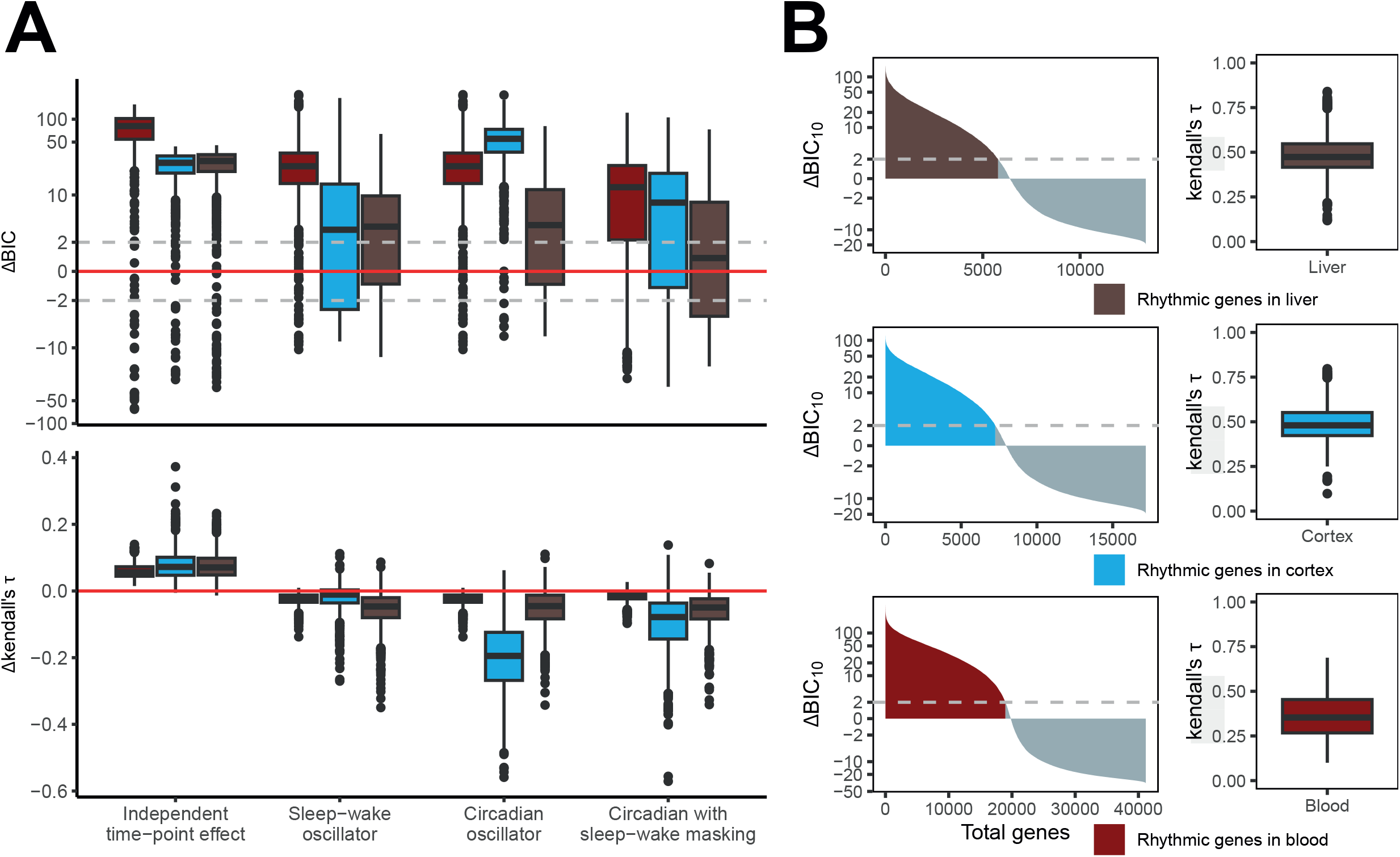
Model performance against alternative hypotheses. **(A)** Our circadian and sleep-wake driven oscillator model (H_1_) versus 4 alternative models (H_A_): i) a linear model with independent time-point effect, ii) a sleep-wake driven oscillator only, iii) a circadian driven oscillator only, iv) a circadian function with an additive effect of sleep-wake (‘masking’; see Results). ∆BIC (upper panels) of H_1_ vs. H_A_ for 1000 rhythmic genes and probes during baseline for blood (red), cortex (blue), and liver (brown). Positive values represent a better fit for H_1_, negative values a better fit for H_A_. Values between -2 and +2 can be considered as low evidence for either model. ∆Kendall’s tau (lower panels) of H_1_ vs. H_A_ shows goodness of fit between models. Negative values support H_1_. **(B)** Detection of rhythmic genes in the entire transcriptome. ∆BIC of H_1_ versus the null hypothesis H_0_ of no rhythmic expression (*yi* = *β*0 +*ε*). Genes and probes with a ∆BIC > 2 are considered to be sleep-wake and/or circadian driven resulting in their rhythmic expression under unperturbed and/or perturbed conditions in liver (top, brown), cortex (middle, blue), and blood (bottom, red). Right panels: goodness of fit (Kendall’s tau) for rhythmic genes in the three tissues. Boxplots depict Kendall’s tau between fitted values and observed value for rhythmic genes in liver, cortex, and blood.

This analysis supports H_1_ as it importantly improved the overall fit, while model complexity did not increase too much over simpler models. Although expression dynamics of individual genes might be fit better with simpler models, the use of a single model for all genes and letting the parameter optimization decide which of the drive is dominant has important advantages as it avoids having to determine the optimal model for each gene. Moreover, using multiple models renders parameter comparison among genes (or for the same genes in different tissues) hard, if not impossible.

### The cortical transcriptome is mainly sleep-wake driven, that of liver and blood mainly circadian

We then applied the H_1_ model to the entire transcriptome to detect, in an unbiased manner, any gene that would be sleep-wake driven and/or circadian driven by contrasting the results to a flat model with a single intercept as null hypothesis (H_0_) where expression variance represents noise. With a ΔBIC>2 as rejection threshold, the model classified a surprisingly large number of genes as rhythmically expressed: 7’246 (42% of 17’185) and 5’785 (43% of 13’373) genes in cortex and liver, respectively, and 18’954 probes (46% of 41’162) in blood (**Fig. 4B**). The high number of rhythmic genes compared to that reported in other studies [1, 2, 50] is likely because the model combines circadian and sleep-wake contributions giving rise to more complex dynamics than can be fitted with simpler sinewave function or time courses with small amplitudes under undisturbed conditions due to opposing contribution of the two forces, such as illustrated with *NCOR1* (**Fig. 3**). Mean goodness of fit as Kendall’s τ for rhythmic genes is high in cortex and liver (∼0.5, **Fig. 4B - right**). Model fit was lower for the human blood dataset compared to that obtained for the mouse datasets both for overall probes as well as for the 1000 rhythmic probes (Δ mean Kendall’s tau: 0.17 and 0.12, respectively) but nevertheless still close to that of the more complex model (**Fig. 4A**).

To assess and visualize the main source of variance for these rhythmically expressed genes, we performed a principal component analysis (PCA; **Fig. 5, Fig. S3**) and projected the model fits in PCA space together with the corresponding circadian and sleep-wake driven responses plotted alongside the PC axes to show their respective contributions for the time segments depicted in the PC plots (**Fig. 5A-D**). In addition, the complete simulated time-course of the responses to *f=f*_*S*_ and *f*_*C*_ for the first two principal components, PC1 and -2, is illustrated underneath each panel for each of the experiments.

**Figure 5:**
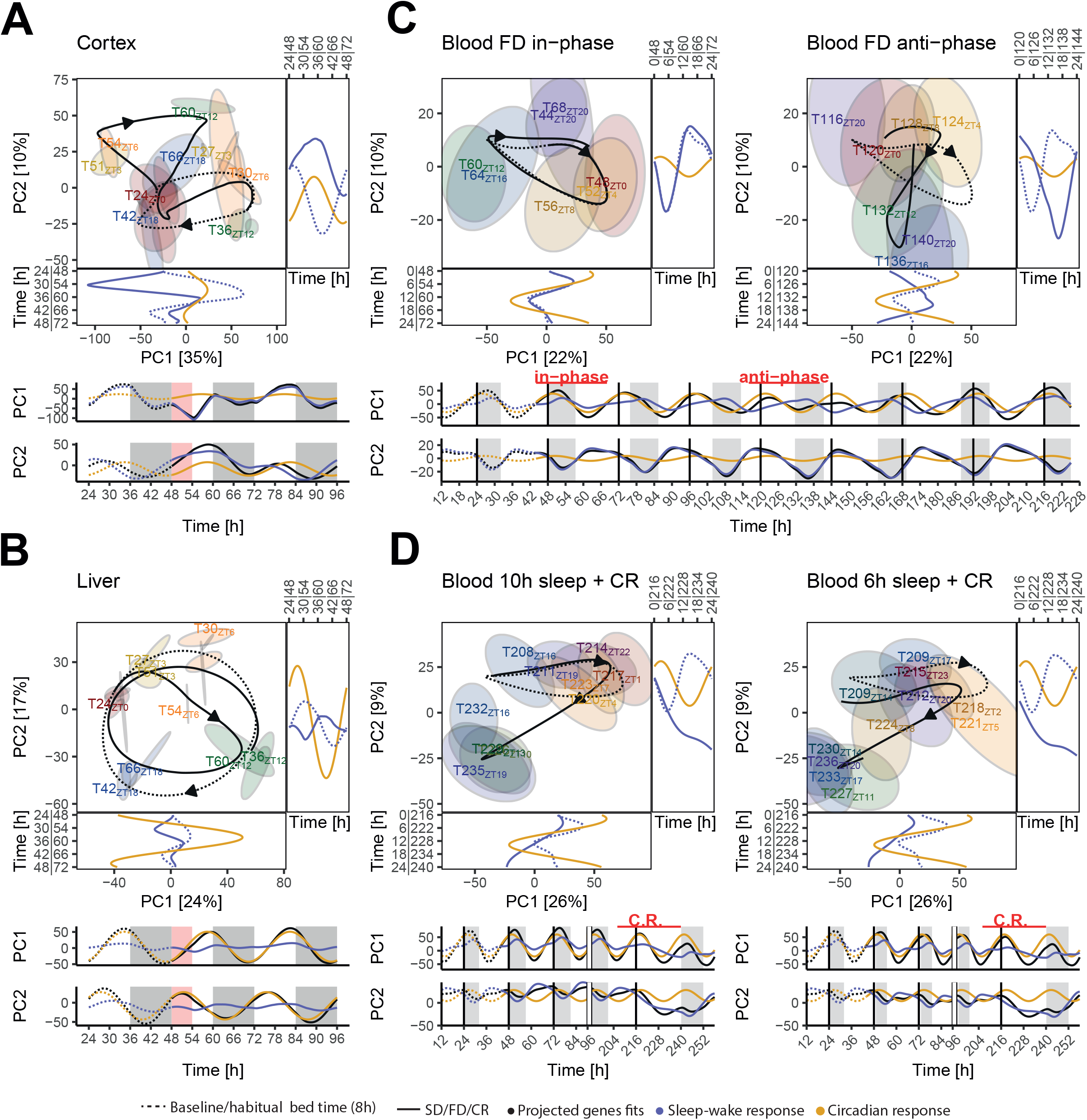
Principal component analysis (PCA) of the rhythmic transcriptomes. **(A)** PCA in the mouse cortex and **(B)** liver during baseline (BSL), sleep deprivation (SD), and recovery (REC), **(C)** in human blood during the Forced Desynchrony (FD) when sleeping ‘in-phase’ (left) and ‘anti-phase’ (right panels), and **(D)** in human blood during the Constant Routine (CR) after the 10h sleep (left) and 6h sleep opportunity (right panels). Variance explained by each PC in brackets. Projected model fits in PCA space during BSL and habitual bedtime as dashed lines, fitted expression during SD + REC, FD, and CR conditions as solid lines. Arrowheads point into the direction of the progression in time. Ellipses delimit 95% confidence intervals of data acquired at each time point. Corresponding circadian (yellow) and sleep-wake (purple line) driven responses are plotted alongside the PC axes. Note double labels at time axes corresponding to the respective times in the experiment for the two conditions (see time courses below). The complete simulated time-course of the circadian and sleep-wake driven responses for PC1 and -2 is illustrated underneath each panel for each of the experiments. Pink and grey boxes indicate the SD and dark periods, respectively, in mice; grey boxes for human experiments the scheduled sleep episodes.

Distinct types of dynamics could be observed in the mouse transcriptomes. In cortex, PC1 displayed a predominant sleep-wake driven response (projected *SWrc* = 0.80) composed of overdamped genes as top contributors, with a large immediate effect of SD and a subsequent quick recovery (**Fig. 5A**), a pattern consistent with that of sleep-wake driven IEGs and the strong chromatin remodeling effect of SD in this tissue [14]. GO analyses identified that these genes are involved in protein folding, RNA regulation, and chromatin organization (**Fig. S3 - Cortex**). PC2, on the other hand, was determined by underdamped genes with large phase-lags 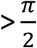 rad) responding to both circadian and sleep-wake drives (*SWrc* = 0.60). The latter drive increased gene expression during SD which continued during the first 6h of recovery (i.e., until ZT12 of the first recovery day; ZT12_REC_ in **Fig. 5A**) although mice were mostly asleep during this period. The model found that this inertia in the response to the SD was a consequence of a weak negative feedback and a large phase-lag. Evidence of such inertia can already be observed in baseline when increases followed the sleep-wake distribution with a similar long delay (start of increase at ZT15, i.e., ca. 3h after spontaneous wake onset at lights-off, until ZT3, **Fig. 5A - lower panel**, dashed blue line for PC2, **Fig. S3 - Cortex**). These genes are involved in neurotransmitter transport/signaling, feeding behavior, phosphatidylinositol dephosphorylation, and fatty acid metabolism.

In liver, the fitted trajectories for the expression of genes contributing to PC1 and -2 followed circular patterns and both PCs showed a large contribution of the circadian response relative to the sleep-wake response (*SWrc* = 0.18 and 0.29, respectively) albeit with different phases (**Fig. 5B, Fig. S3 - Liver**). SD decreased the amplitude of PC2 (ZT6_SD_) and was followed by an amplitude reduction 12h later (ZT18_REC_). PC2 shows an enrichment for genes involved in androgen receptor signaling and, similar to PC1 genes in cortex, in protein folding. PC1 genes in liver were left largely unperturbed by SD and were enriched for genes implicated in GTPase activity. The response dynamics for transcripts contributing to PC2 in liver and cortex highlight a novel and slower type of sleep-wake driven response requiring more time to change mRNA levels compared to the fast IEG (and delta-power) -like response observed for PC1 in cortex.

For the human blood transcriptome, PCAs of the FD ‘in-phase’ and ‘anti-phase’ conditions (**Fig. 5C**) and the CRs after the 10- and 6h sleep-opportunity conditions (**Fig. 5D**) were plotted separately for better visualization. The predicted expression dynamics during habitual bedtime (24h sleep-wake cycle with 7.5h sleep) was used as a ‘baseline’ reference (dashed lines in **Fig. 5C-D**). The expression dynamics fitted to the FD ‘in-phase’ condition were at first indistinguishable from the predicted baseline dynamics (**Fig 5C - left**) and deviations appeared only after ZT8 (i.e., 8h after the onset of the scheduled sleep episode) when under baseline subjects woke up, while under the FD condition sleep was scheduled to last an additional 1.3h. As PC2 was mostly sleep-wake driven (*SWrc* = 0.80), due to the longer sleep periods and the longer wake periods of the 28h day compared to the 24h day, the amplitudes of the rhythmic probes contributing to PC2 gradually increased over the initial 4 days of the FD to a new steady-state (bottom time-course in **Fig. 5C**). Thus, by the time the ‘anti-phase’ condition was reached, PC2 showed a strong amplitude increase (**Fig. 5C - right**). Top contributors to PC2 were mostly underdamped probes with weak negative feedback such as *NCOR1*. The corresponding genes were found to be involved in B-cell activation and phosphatidylinositol dephosphorylation, the latter confirming the PC2 pathway found in the mouse cortex (**Fig. S3 - FD**).

PC1 was more circadian than sleep-wake driven (*SWrc* = 0.36). Probes contributing to PC1 were enriched for genes involved in translation and mitochondrial regulation (**Fig. S3 - FD**). PC1’s overall amplitude reduced during the ‘anti-phase’ condition when sleep-wake and circadian responses opposed each other (**Fig. 5C - bottom panel**). The model predicted an even more prominent amplitude reduction during the 28h day following ‘anti-phase’. As we had access to sleep-wake data throughout the 10-day FD protocol (**Fig. 1B**), we could simulate expression dynamics when subjects returned to being ‘in-phase’ again 3 days later (i.e., the last day of the FD) and found that the amplitudes of both PC1 and -2 were larger compared to the ‘in-phase’ condition at the beginning of the FD (**Fig. 5C - bottom panel**). To conclude, the model predicts that over the course of the FD protocol expression dynamics change and that the two ‘in-phase’ conditions will importantly differ transcriptionally.

PCA for the second human transcriptome experiment showed the large effect of the 40h wakefulness during the two CRs importantly amplifying the sleep-wake response contributing to PC2 (*SWrc* = 0.56), as already illustrated for *NCOR1* (**Fig. 3**). The preceding 7 days of restricted sleep changed the initial condition of the CR compared to that of the control condition (6 vs. 10h sleep opportunity) again affecting mostly PC2, the trajectory of which was downshifted during the CR (**Fig. 5D - left vs. right panel**). This could also be observed at the level of the data where ellipses, denoting the 95% CI of mean gene expression, were all slightly lower after the restricted sleep condition. The model predicted that the lowering of PC2 already occurred on the 2^nd^ day of the sleep restriction protocol (**Fig. 5D - bottom-right panel**). In contrast, for the 10h-sleep-opportunity condition the model found an increase in PC2 over the first days of the protocol compared to baseline before slowly decreasing again reaching baseline level prior to the start of the CR. This increase and subsequent decrease can be attributed to the initial increase in mean total sleep time in the first days of the protocol (9.4h on the first day) that then reverted to baseline levels (7.7h on the last day; **Fig. 1C - bottom-left panel**). Like FD, PC1 is enriched for translational regulation, and PC2 for cell division and protein lipidation (**Fig. S3 - CR**).

Comparing the PCA across species and tissues showed some surprising similarities considering they were computed independently. Although the relative contribution of the sleep-wake response and the circadian response varied among tissues, PC1 showed a mostly in-phase relationship between the two responses during baseline for all datasets (**Fig. S3**; φ(*Cr* − *SWr*) black line represents the in-phase relationship), while for PC2 their phases importantly differed. Accordingly, the mean amplitudes of genes in PC1 are larger than that of PC2 genes in all tissues (Cortex: 0.25 vs. 0.11, Liver: 0.66 vs. 0.48, Blood: 0.22 vs. 0.10, p-values < 1e-9). This did, however, not translate into common genes contributing to each of two PCs across datasets. Only PC1 of the FD and CR experiments had a strong concordance of contributing genes. This suggests that the biological processes that are sleep-wake and circadian driven differ across tissues, with most genes displaying an in-phase relationship, and a smaller proportion of the transcriptome with an anti-phasic relationship. The expression of the latter class of genes represented by PC2 appears more prone to long-term deviations from baseline upon perturbation of sleep with a larger sleep-wake contribution (except cortex) and lower *γ* (**Fig. S3**) which increases the time needed to again reach baseline dynamics (ττ, Eq. 4).

As the PCA reports only on those transcripts contributing most to the overall variance, we assessed the *SWrc* values for the complete rhythmic transcriptome. As already indicated by its PC1, the model found that cortical gene transcription was more sleep-wake driven than in liver and in blood, with similar *SWrc* values obtained in the latter two tissues (mean *SWrc*: 0.62, 0.37, and 0.40 for cortex, liver, and blood respectively; **Fig. 6A**). In cortex 67% of rhythmic genes were underdamped (ζ < 1), while 85% and 89% of genes in liver and blood were underdamped. Although mostly underdamped, the analysis of rhythmic genes in blood revealed a conspicuous cluster of overdamped transcripts (ζ > 2) that were mostly circadian driven (*SWrc* < 0.5). GO analysis of this overdamped cluster revealed an enrichment for genes involved in acetylcholine receptor binding that were strongly circadian driven (*SWrc* < 0.25) and, for the remaining transcripts in this cluster (*SWrc* > 0.25), genes involved in signaling adaptor activity and dopamine receptor binding. We compared our results in blood with the classification made by Archer and colleagues based on the FD transcriptome results using an additive model with the free-running circadian melatonin rhythm and the enforced 28h sleep-wake cycle as factors [23] (**Fig. S4**). As expected, probes originally classified as changing in-phase with melatonin have, in our model, a low *SWrc* (mean = 0.25) and probes classified as in-phase with the sleep-wake cycle have a high *SWrc* (mean = 0.60). There were, however, some noticeable exceptions, such as *SERPINB9* which was categorized as in-phase with melatonin, suggesting an important circadian influence, whereas our model found its expression to be strongly sleep-wake driven (*SWrc* = 0.82; **Fig. S4**). Because of its long time-constant (τ = 42.0h), *SERPINB9* expression was only slightly shifted at the time sleep occurred in anti-phase with the melatonin rhythm and ca. 3 additional days of sleeping in anti-phase (i.e., 2 * τ) would have been required to observe a more complete shift of *SERPINB9* expression relative to the melatonin rhythm such that it again realigns with the sleep-wake distribution. Consistent with the prediction of a sleep-wake driven oscillatory (underdamped) dynamics with an amplitude reduction by extended wakefulness, blood *SERPINB9* expression was found to be down-regulated after SD [51] and rhythmic in an independent CR experiment [52].

**Figure 6:**
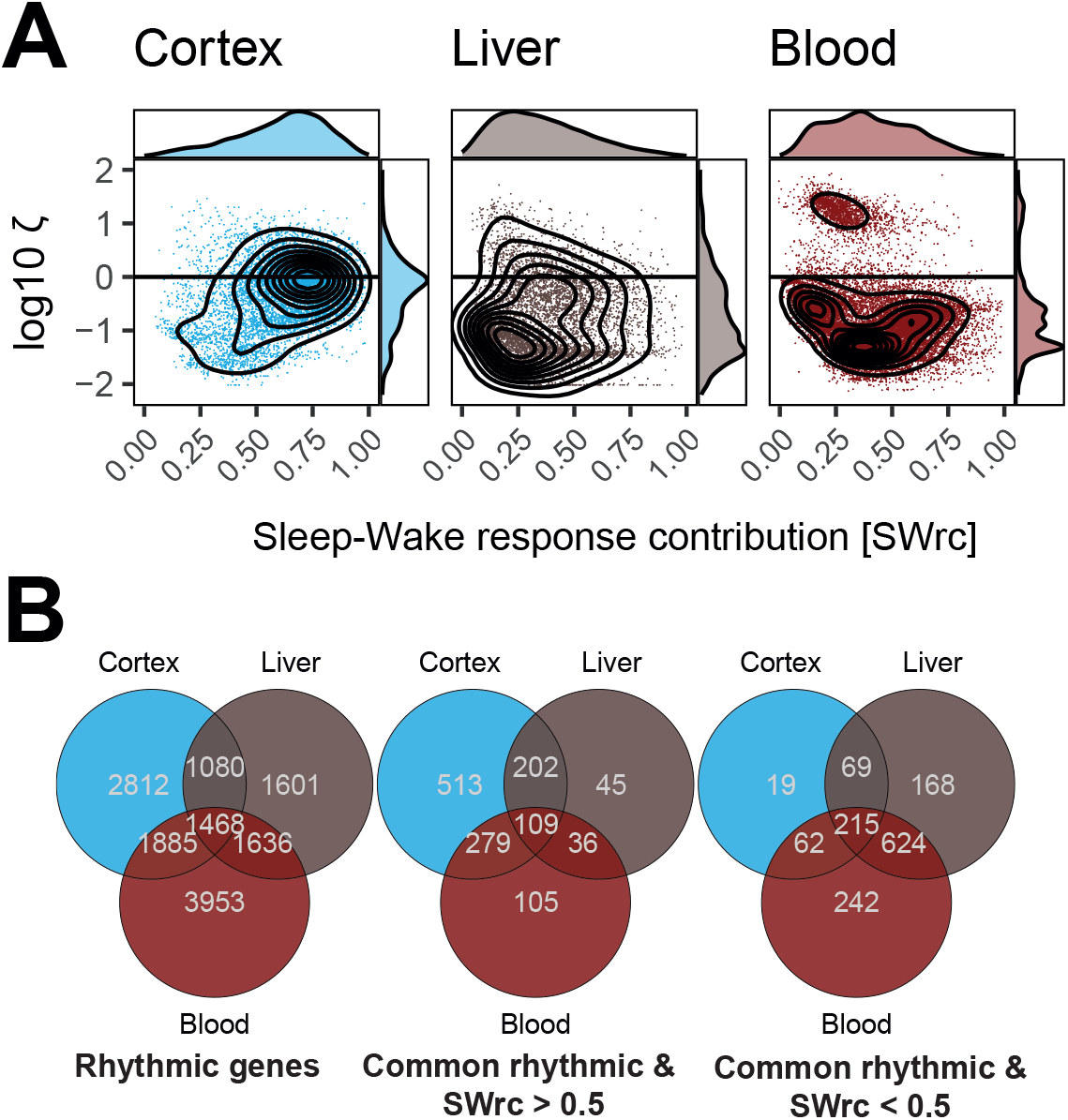
Relative contribution of circadian and sleep-wake driven responses to gene expression. **(A)** Relative sleep-wake response contribution (*SWrc*, see Results) versus damping ratio (ζ) for all rhythmic genes in cortex (blue), liver (brown), and blood (red dots). Black lines represent 2D gene density. **(B)** Venn diagrams of all rhythmic genes (left) and the 1425 rhythmic genes common among the three tissues: Sleep-wake driven (*SWrc* > 0.5, middle) and circadian driven (*SWrc* < 0.5, right panel) genes in mouse cortex and liver and human blood.

Of all genes found to be rhythmic across the datasets (14’435), only 10% (1468) were rhythmic in all 3 tissues (**Fig. 6B**). This strong tissue specificity of gene rhythmicity has already been noted in other species [53]. We then compared the *SWrc* of these 1468 shared rhythmic genes but did not find any correlations between tissues (cortex *vs*. liver pearson correlation: 0.08, cortex *vs*. blood: -0.001, liver *vs*. blood: 0.003) indicating that the cause of rhythmicity (circadian *vs*. sleep-wake driven) was not shared. Nevertheless, most of the few genes found to be sleep-wake driven in liver (*SWrc* > 0.5) were also sleep-wake driven in cortex (311; 79% of 392), and the circadian-driven genes in cortex (*SWrc* < 0.5) were also circadian driven in liver (78%; **Fig. 6B**).

Among the 1468 common rhythmic genes, only 109 had *SWrc* values above 0.5 in all 3 tissues (**Fig. S6**), with *Ndufs1* as the gene with the highest average *SWrc* (0.82). *Ndufs1* is a mitochondrial gene involved in reactive oxygen metabolism and was previously found as a biomarker for short-sleep duration [54]. Interestingly, the most circadian driven gene among the 1468, *Sod2* (average *SWrc* = 0.10), is also a mitochondrial gene involved in reactive oxygen metabolism. We found that the top-most enriched biological process for the 1468 genes rhythmic in all tissues was protein folding (**Fig. S5**). Protein folding was also found as the most enriched biological process for the 109 sleep-wake driven rhythmic genes shared among the three tissues. Conversely, 215 genes had *SWrc* values below 0.5 in all 3 tissues. These common circadian-driven genes were involved in Protein kinase B signaling. Phosphatidylinositol 3 kinase signaling appeared as the 3^rd^ most significantly enriched GO term, which is interesting as genes contributing to PC2 in mouse cortex and human blood (**Fig. 5**) were enriched for genes involved in the dephosphorylation of phosphatidylinositols, which have been associated with sleep [55-57].

It should be noted that while we considered genes as sleep-wake driven or circadian driven using a *SWrc* cut-off of 0.5, the drive that contributes less still affects gene expression dynamics. For only less than 3% of each of the transcriptomes, genes could be labeled as either entirely sleep-wake driven or entirely circadian driven (*SWrc* > 0.95 or < 0.05). Therefore, for most transcripts both drives need to be considered when studying rhythmic gene expression.

### Sleep deprivation desynchronizes the tissue transcriptome

Although central and tissue rhythms in gene expression are generally associated with clock genes implicated in the TTFL, clock genes did not feature among the top circadian driven genes. We therefore took a closer look at the expression dynamics of 15 core clock genes (**Fig. 7A**). Expression of 11 out of the 12 clock genes that were rhythmically expressed in the cortex showed a mainly sleep-wake driven response (*SWrc* > 0.5). In contrast, in liver and blood, most clock genes were found to be circadian driven (0 and 1 out of 13, respectively; *SWrc* < 0.5). In cortex, *Clock* is the strongest sleep-wake driven clock gene (*SWrc* = 0.84) and among the top 11% most sleep-wake driven genes in this tissue but is mostly circadian driven in liver (*SWrc*: 0.19; **Fig. 2C**) and blood (*SWrc* = 0.26).

**Figure 7:**
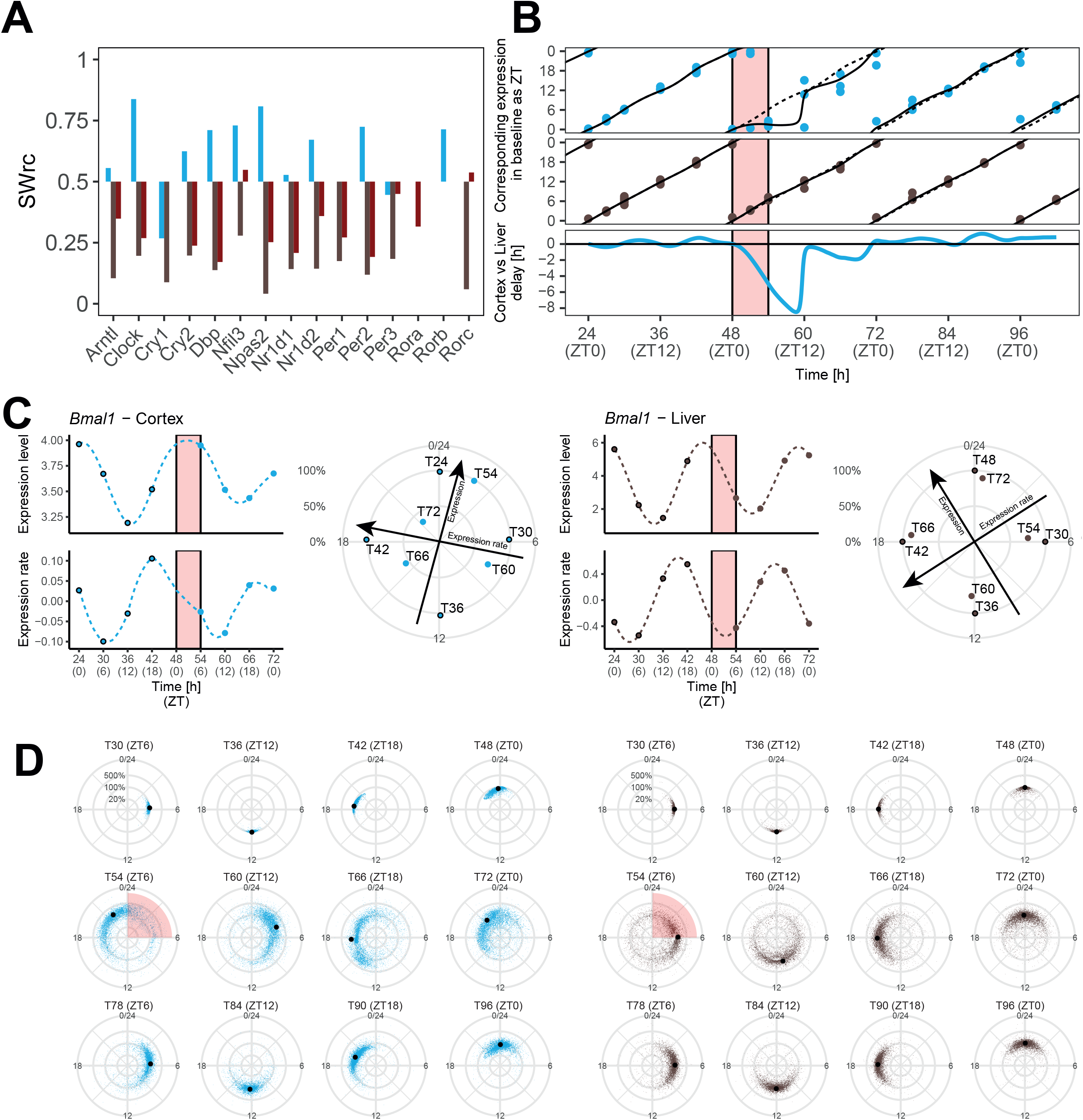
Sleep deprivation (SD) changes timing of gene expression within and between tissues. **(A)** Sleep-wake response contribution (*SWrc*) for clock-gene expression in mouse cortex (blue) and liver (brown) and blood (red) in humans. In blood, mean *SWrc* was estimated from the probes of the same clock genes. **(B)** Fitted and predicted local biological time in cortex and liver based on clock-gene expression. The tissue’s local time [expressed as zeitgeber time (ZT) in baseline; ZT0/24, -3, -6, -12, and -18] was fitted using baseline clock-gene expression with an elastic net model (see Results). Local time is then predicted for gene expression during SD (T51_ZT3_, T54_ZT6_) and subsequent recovery (REC, i.e., T60_ZT12_, ZT66_ZT18_, and T72_ZT0_). Projected fits based on our oscillator model as dashed (baseline) and solid (response to SD) lines. Lower graph depicts the cortex-liver tissue differences in predicted ZT. **(C)** Estimated relative phase and amplitude of *Bmal1* from expression level and expression rate of the model. Baseline points T24/T48_ZT0_, T30_ZT6_, T36_ZT12_, and T42_ZT18_ are fitted to a 24h clock. Time on the horixontal-axes are given both in time-of-experiment and ZT (in parentheses). **(D)** Relative phase and amplitude individually fitted (upper row panel) and predicted (middle/lower panel) for the expression of all rhythmic genes in cortex (left, blue dots) and liver (right, brown dots). Larger black dots represent ‘point-of-gravity’ of level and rate of expression of all genes.

While clock genes in the SCN are involved in timekeeping, their role may be more diverse in tissues peripheral to the SCN [45, 58]. Because clock genes are sleep-wake driven in the cortex and circadian driven in liver, sleep perturbation may alter inter-tissue synchrony and clock-gene related processes like metabolism [59]. To assess tissue differences in cellular timing, we fitted clock-gene expression in cortex and liver to a 24h clock corresponding to the tissue’s zeitgeber time (ZT) in baseline (**Fig. 7B - dashed line**) using a multivariate regression model with elastic net regularization [60]. We observed that during the SD and the subsequent 5h of recovery (corresponding to ZT0-11 in baseline) cortical local time no longer followed ZT and that the expression dynamics of clock genes was halted at a state corresponding to ZT0-2 during baseline (**Fig. 7B - solid line**). In contrast, in the liver, circadian time progressed undisturbed resulting in an important desynchronization between the two tissues with a maximum cortex-to-liver delay of 8h reached 5h after the end of the SD (**Fig. 7B - bottom**).

As the cortical transcriptome, including most clock genes, is mostly sleep-wake driven, zeitgeber time (or circadian time defined by phase markers of the central circadian clock) has little significance in this tissue. That zeitgeber time estimated by the expression of clock genes was maintained at ZT0-2 for 11 consecutive hours does therefore not indicate that the circadian clock stopped but simply results from the SD keeping waking levels high for 6 additional hours following the baseline dark period when animals were mostly awake spontaneously. The limited use of clock genes as biomarkers of circadian time in tissues peripheral to the SCN under conditions of altered sleep-wake distributions has already been suggested previously [61].

The SD causes the cortex and liver transcriptomes to desynchronize as tissue oscillators differ in their overall response to sleep-wake state (**Fig. 6A**). Similarly, within each tissue, genes revealed a wide range of responses (**Fig. 6A**) implying that SD also changes intra-tissue synchronicity. To examine this, we performed a similar analysis as above, where the baseline timing of expression is estimated independently for each gene based on its expression level and expression rate predicted by our model. The baseline time points ZT0, 6, 12, and 18 were mapped to zeitgeber time and time points after the start of the SD plotted according to baseline time considering expression level and expression rate (**Fig. 7C**). In this representation, the distance from the center reflects a relative amplitude change (100% = baseline) and an angular change between corresponding ZT points before (baseline) and after SD (ZT_SD_ and ZT_REC_) can be viewed as a phase change. In the figure each dot represents one gene, and the ‘point of gravity’ of all genes is represented with a black circle. As expression level and expression rate in baseline could not be mapped perfectly to a 24h clock, we observed small scattering around the points of gravity at the four time points (**Fig. 7D - upper panels**). Rhythmic genes which could not be readily mapped to a 24h clock (because their baseline time course deviated too much from a sinewave like dynamic; R^2^ < 0.6, see Methods), and thus scattered too much, were excluded from this analysis (9 and 4% of all rhythmic genes in cortex and liver, respectively). SD caused extensive scattering of gene timing in both tissues which lasted for more than 24h (**Fig. 7D - middle and lower panels**), indicating that the phase relationship among genes is largely altered by SD. Despite this increased scattering, the point of gravity in liver still closely followed baseline timing. In contrast, in cortex overall timing was greatly impacted with points of gravity deviating from those observed in baseline by ca. 8h at ZT6 and -12. It thus appears that the SD-induced changes in the cortical timing of expression level and expression change observed of clock genes (**Fig. 7B**) apply to the entire rhythmic transcriptome in this tissue. On the second recovery day, scattering of timing remained larger than in baseline in both tissues, suggesting that the expression of many genes was still perturbed although in cortex the location of the points of gravity suggest that overall, the timing had reverted to that of baseline (**Fig. 7D - lower panels**).

### Does recovery sleep accelerate transcriptome recovery?

We previously reported that the expression dynamics of a large number of genes affected by SD still deviated from baseline long after the sleep-wake distribution and EEG activity had reverted to baseline, i.e., beyond the first 18h after the SD ended [14] (see **Fig. 7D**). Using our model prediction, we further investigated the ‘recovery’ dynamics for the rhythmic transcripts affected by SD, i.e., those with a fold-change effect size > 1 [z-score] at any time-point during the 48h after SD. We first determined how the fold-change in expression reached at the end of SD (ZT6_SD_) related to the time required for expression to again reach equilibrium, i.e., the time constant, τ (Eq. 4). Perhaps counter-intuitively, we found that, in general, genes for which the expression was affected the most at the end of the SD had the shortest time-constants (**Fig. 8A**). More genes displayed such a strong-and-fast response in cortex than in liver where the initial responses tended to be smaller but longer lasting (**Fig. 8A**).

**Figure 8:**
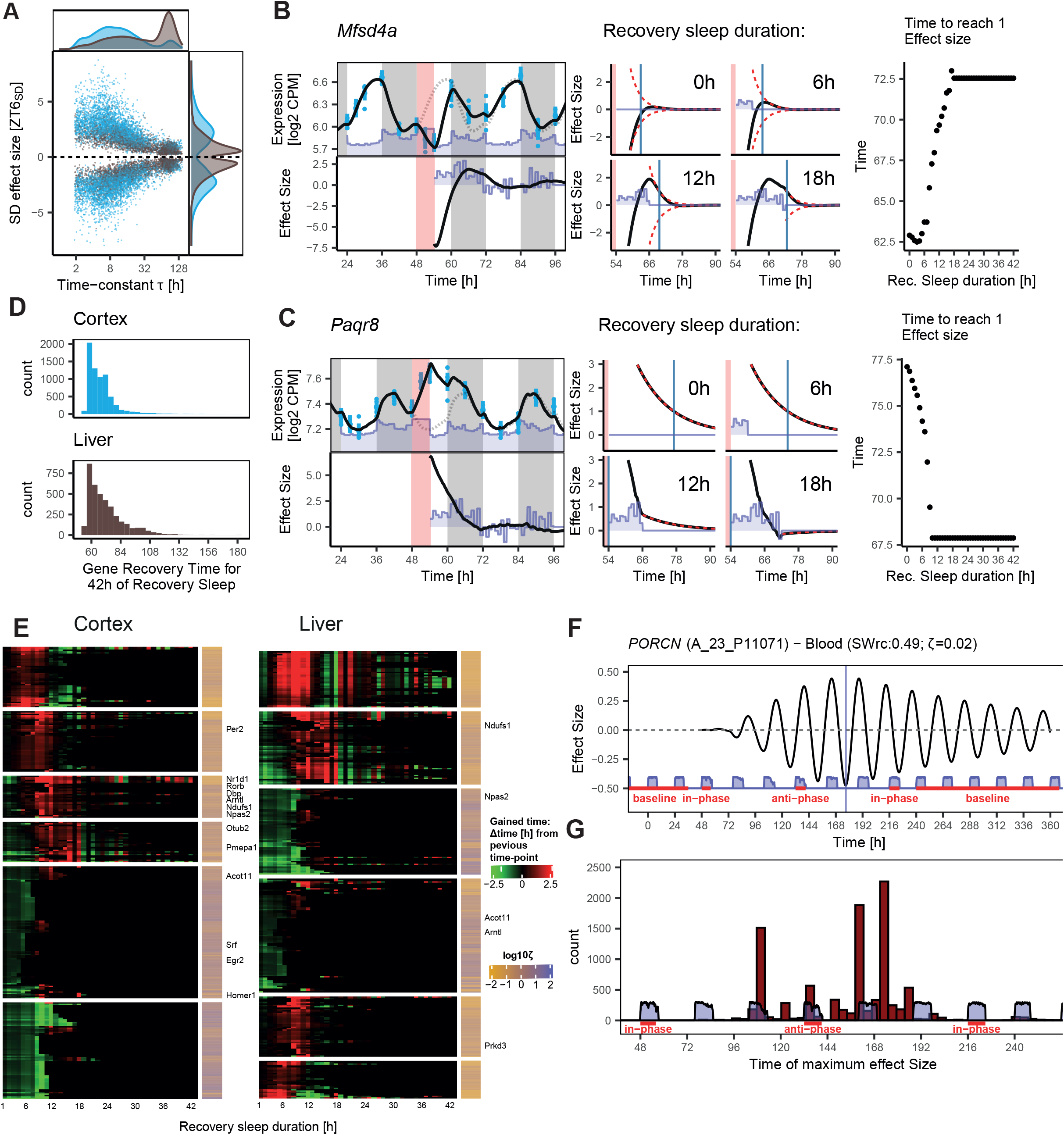
Responses to recovery sleep. **(A)** Effect-size of differential gene expression at the end of sleep-deprivation (SD; ZT6_SD_ vs. ZT6 in baseline) versus the model-derived recovery time-constant τ in mouse liver (brown) and cortex (blue) for all rhythmic genes with a sleep-wake driven contribution (*SWrc* > 0.25). Relative distributions for τ and effect size plotted along their respective axes. **(B)** Left panel: *Mfsd4a* expression (blue bars, 95% ci), its model fit (solid black line; dotted line replots baseline fit), and sleep-wake distribution (purple area; upper graph), with recovery vs. baseline effect-size (black line) after SD and hourly values of sleep gain during recovery (purple area; lower graph). Center panels: Effect-size (black lines) when 0-, 6-, 12-, or 18h of the actual recovery sleep recording (as opposed to baseline sleep) was used for predicting gene expression after SD. Purple area indicates sleep gain included in each of the 4 simulations. Dashed red lines are the exponential parts of the oscillator solution when using only baseline sleep after SD (0h recovery sleep; also see **Fig. 2B - left panels**). Blue vertical line marks the time-point at which the exponential part reaches an effect size of +1.0 or -1.0, which in subsequent analyses is considered the time at which gene expression has recovered. Right panels: Time point of gene recovery when including 0- to 42h of recovery sleep. **(C)** as B but for *Paqr8*. **(D)** Histogram of gene recovery time-points for all rhythmic genes with a *SWrc* > 0.25 in cortex (upper, blue) and liver (lower panel, brown) using the actual (42h) recovery sleep. **(E)** Gain in gene recovery time for all genes in D in cortex (left) and liver (right). Analyses as in right-hand panels of B and C but here the differences from one time-point to its preceding time-point are plotted. As more sleep recovery recording was included in the simulation, genes either advanced (green) or delayed (red) their recovery time. Data were filtered to show only genes with a minimum of 1h advance or delay. **(F)** Effect size for differential *PORCN* expression (FD vs. baseline) for the expression simulated during the entire FD protocol and for 10 repetitions of baseline sleep-wake patterns under 24h days after the second in-phase condition. Vertical blue line indicates the time when maximum effect size was reached (Time = 177h). **(G)** Time of maximum effect size modeled in the FD protocol for all rhythmic genes.

The immediate SD effect and τ alone were, however, insufficient to account for the large variability among genes and tissues in the time required for gene expression to recover. One factor that could play a role is the extra sleep gained during recovery, which could be viewed as a second perturbation shortening or lengthening the duration for a gene to return to its baseline rhythmicity. In fact, τ correctly estimates the time to return to baseline equilibrium only if mice do not alter their sleep-wake behavior after SD, i.e., sleep as during baseline. To evaluate the effect of recovery sleep on transcriptome recovery, we simulated gene expression in mice that do not (referred to as ‘0h recovery sleep’) or partially compensate for sleep loss by incrementally (hour-by-hour) replacing the subsequent sleep-wake distribution by their ZT-matched baseline sleep. We illustrate this analyses with the simulated expression of *Mfsd4a* and *Paqr8* with either 0-, 6-, 12-, or 18h of recovery sleep (**Fig. 8B**,**C - middle panels**). We took these two genes because their cortical response to recovery sleep was opposite while both tended to be sleep-wake driven (*SWrc* = 0.80 and 0.52) and showed a comparable large effect size after SD (-7.0 and +6.5, respectively at ZT6_SD_; **Fig. 8B**,**C - left panels**). Moreover, *Mfsd4a* and *Paqr8* were under- and overdamped, respectively (ζ = 0.76 and 3.17). From the time point in the simulation when the actual recovery sleep was replaced with baseline sleep, the fold-change of underdamped genes (such as *Mfsd4a*) can be viewed as an underdamped oscillator relaxing back to equilibrium with its amplitude decaying exponentially with a time-constant τ (red dashed lines in **Fig. 8B – middle panels**; compare to *Gene A* in **Fig. 2B – left panel**). For overdamped genes (such as *Paqr8*), the reduction in the fold-change follows a simple exponential decay (red dashed lines in **Fig. 8C – middle panels**; see *Gene Y* in **Fig. 2B – left panel**). We then calculated the time required for the exponential decay part describing the recovery of gene expression to reach an effect-size of < 1 and considered gene expression to have recovered at this time-point. We estimated that 50% of all genes affected by SD ‘recovered’ within 12 and 13h, and an additional 17 and 12% after 18h of recovery, for cortex and liver, respectively (**Fig. 8D**). This implies that at the time sleep and EEG phenotypes no longer differed from baseline, the expression of 32 to 37% of genes still had not recovered. Using the baseline sleep-wake data instead of the actual sleep-wake recovery data accelerated the recovery of *Mfsd4a* expression by approximately 10h, while it delayed *Paqr8*’s recovery by a similar duration (**Fig. 8B,C - middle panels**). Or, in other words, as more recovery sleep was included, time of recovery increased for *Mfsd4a* from 62.h to 72.5h when 10h of recovery sleep was included, and decreased for *Paqr8* expression (from 77.4 to 67.9h) with 18h of recovery sleep (**Fig. 8B,C - right panels**). In general, overdamped (log_10_ ζ > 0) genes, such as *Paqr8*, seemed to benefit from sleeping more (**Fig. 8E**, green-black sequence, with green indicating that including 1h of recovery sleep accelerated gene recovery), whereas most genes with an oscillatory component (i.e., underdamped, like *Mfsd4a*) delayed their recovery time as more of the actual recovery sleep was being used for the simulation (**Fig 8E**, red-black sequence). We also observed more complex responses where recovery sleep initially decreases and subsequently increases recovery time (**Fig. 8E**, green-red-black sequence). The opposite sequence could also be observed (**Fig. 8E**, red-green-black sequence). Clustering the response of all genes revealed the presence of 6 types of responses (**Fig. 8E**). In the cortex recovery sleep delays gene recovery time for most of the clock genes. In contrast, several IEGs genes like *Homer1, Srf*, and *Egr2* and others like *Acot11* take advantage of the extra sleep after sleep deprivation to recover faster. Sleep-wake driven genes like *Ndufs1*, and top contributors to first two PCs (**Fig 5A,B**), such as *Otub2* (cortex PC1), *Pmepa1* (cortex PC2), and *Prkd3* (liver PC2), showed gene recovery times that mostly delay when allowing recovery sleep for 6-18h.

The previous analysis emphasized that transcriptome recovery outlasts sleep-wake recovery, and, in addition, that not only a lack of sleep (SD) but also extra time-spent-asleep (recovery sleep) can delay attaining baseline gene-expression dynamics. Given these insights, we explored the transcriptome dynamics during the FD protocol during which subjects recover from transitioning from sleeping in-phase to anti-phase and back again by calculating the gene effect size of the predicted differential expression to corresponding baseline ZT time points. For each gene, we calculated the time-point at which the effect-size was highest. For example, for *PORCN*, a gene with a large effect size (top 2%) and extreme long time constant of recovery (τ = 160h), maximum effect size was reached at time 177h (**Fig. 8F**). The model predicted that for most genes, the largest effect sizes occurred around that time (144-192h), i.e., during the 28h day that followed the anti-phase condition (**Fig. 8G**; Day 7-8 of the protocol, **Fig. 2B**). Such delayed response is reminiscent of the delayed gene-expression responses observed in mice after SD. The model also predicted that genes can still deviate from their baseline dynamics when sleep occurred again in-phase such as, e.g., PORCN (**Fig. 8F**) which might, however, be difficult to demonstrate statistically because of the small, predicted effect size.

## Conclusions

We have presented a mathematical framework that can describe and predict rhythmic gene expression in brain and body tissues peripheral to the SCN. The model integrates and quantifies the contributions of circadian and sleep-wake state related factors and their interaction acting on the daily changes in mRNA levels. The respective contributions of these factors were represented as two drives that each alter the acceleration of the ongoing changes in gene expression within the cells of the tissue. The model was able to capture the often complex and sometimes counterintuitive relationships between sleep-wake interventions, circadian time, and gene expression in cortex and liver in mice and in blood in humans. One strength of the model is that it accommodates within one and the same mathematical framework a variety of expression dynamics. This has the important advantage that parameter optimization will decide with which type of dynamics each gene responds to the exerted drives and which of the two drives is dominant. The model successfully captured changes in gene expression under a number of experimental conditions that altered sleep-wake timing relative to circadian timing, while keeping the number of free parameters low. Applying the model to mouse and human time-course transcriptome data yielded several new insights that are summarized below. Our work shows that the daily or circadian changes in *in vivo* gene expression can only be understood when the contribution of sleep-wake history are taken into account. We believe this framework can also be useful to describe and predict the daily changes in other physiological variables and behaviors.

### An alternative response dynamics to extended wakefulness

The effects of sleep loss on neurophysiology, performance, and behavior are often put into the context of the two-process model of sleep regulation with a sleep-wake driven process increasing and decreasing during wakefulness and sleep, respectively, according to exponential (saturating) functions. This process was originally modelled on the dynamics of the sleep-wake driven changes in EEG delta power [62] and, as we showed here (and elsewhere [14, 15, 45], this type of dynamics captured well the changes in the cortical mRNA levels of activity-induced immediate-early genes (IEGs) characterized as overdamped in the model. Accordingly, expression of this class of genes responded to sleep deprivation with a large immediate increase, to then quickly decrease during sleep reaching baseline levels within 7h, i.e., the median time of gene recovery for the 2037 overdamped sleep-wake driven genes. This steep decline following sleep deprivation, which drove gene expression away from a lower asymptote, is typical of an exponential decreasing function and of IEG expression dynamics. Therefore, although it does require the animal to sleep, its fast recovery dynamics is largely independent of rebound sleep, i.e., the increase in time-spent-asleep after sleep deprivation beyond that observed in baseline.

Our current analyses showed, however, that most of the predominantly sleep-wake driven transcripts did not behave like EEG delta power and followed a response dynamic characterized with a small response at the end of sleep deprivation, a slow recovery (16.9h median gene recovery in cortex for the 3469 sleep-wake driven and underdamped genes) and a larger variety of expression patterns. Among these patterns, some genes showed a marked inertia in the response to altered timing of sleep-wake state, with differences in gene expression becoming evident only after some delay. This explains why these transcripts have gone unnoticed in experimental designs that aimed at finding the molecular correlates of the process reflected by EEG delta power and therefore only focused on the immediate effects of sleep loss. The genes following these slower sleep-wake state driven dynamics might be implicated in the homeostatic regulation of time-spent-asleep, which differs from that of EEG delta power in that it has slower dynamics and becomes evident only after EEG delta power has reverted to baseline.

### Unexpected effects of recovery sleep on transcriptome ‘recovery’

Our analyses showed that deviations from the baseline sleep-wake state time-course altered gene expression patterns. Perhaps counterintuitively, these deviations included rebound sleep subsequent to sleep deprivation, which is generally considered to help restore homeostatic balance. Rebound sleep especially affected the genes that responded with slower response dynamics and had an oscillatory component (i.e., underdamped) by delaying their recovery. The combination of the inertia to respond to enforced waking and their sensitivity to rebound sleep resulted in a flattening of rhythm amplitude that lasted well beyond the sleep-wake distribution and EEG activity had reverted to baseline. The cortical expression pattern of most of the core clock genes followed this pattern.

We have used the term gene expression ‘recovery’ as shorthand for describing the time it took to again reach the baseline time course without knowing whether the transcripts indeed play a role in the recovery processes associated with sleep. Among the pathways enriched for sleep-wake driven genes, we found pathways related to chaperon-mediated protein folding in cortex, liver, and blood. Chaperons were found to be associated with consolidated sleep [63] and reduced ER (endoplasmic reticulum) stress. Many lipidic pathways were also enriched for sleep-wake driven genes in both cortex and blood, like those involved in cholesterol/lipid regulation as well as their proportions and spatial arrangement in the cellular membrane.

### Circadian timing and the effects of sleep loss

Our analyses showed that sleep deprivation in the mouse caused a long-term change in the phase relationship among genes within and between tissues. Consistent with more genes being sleep-wake driven in cortex than in liver, sleep deprivation impacted overall timing in cortex to a much larger extent, resulting in a large difference in circadian timing between the two tissues, which amounted to an estimated 8-hour phase delay, 5 hours after the end of the sleep deprivation. The phase differences were observed at the level of the whole transcriptome as well as among clock genes. In cortex, but not in liver, all but one of the clock genes were affected by sleep-wake state with *Clock* and *Npas2* expression, the two transcription factors forming the positive arm of the circadian TTFL, responding, like IEGs, almost exclusively to the sleep-wake time course over the 4-day experiment. This tissue difference in the behavior of clock genes might not surprise given the fact that sleep-wake state is tightly coupled to metabolic activity in the cortex and less so in liver. The clock-gene circuitry in the cortex might thus be used to track and predict time-spent-awake instead of setting circadian time. Accordingly, clock genes in the cortex are of little significance as phase markers of the central circadian clock, as was already suggested by others for other tissues peripheral to the SCN (Dijk and Duffy 2020). To further investigate the relationship between the tissue’s activity and clock gene dynamics, one could, e.g., change (metabolic) activity of the liver specifically without affecting sleep-wake state. We predict that the expression dynamics of clock genes in the liver would become less circadian and more ‘cortex’ like.

## Methods

### Mouse datasets

Mouse transcriptome dataset is available on GEO (TBD). Experimental details are available [12, 14]. The following methods are a summary.

### Animals

62 male mice C57Bl/6J were purchased at Charles River France for RNA-sequencing of cortical and liver tissues. 12 male mice C57Bl/6J were purchased from the University of Tennessee Health Science Center (Memphis, TN, United States of America) for EEG/EMG recording. Both sets of mice underwent same housing condition: mice were acclimated to our facility for 2-4 weeks prior experimental procedure. Mice were kept under 12h light -12h dark conditions. Both experimental procedures were performed at the age of 10-12 weeks and approved by the veterinary authorities of the state of Vaud (SCAV). No additional animal experiments were performed for this publication.

### Sleep deprivation

Sleep deprivation was performed by gentle handling [64] for 6h at light onset (zeitgeber time ZT0-6).

### EEG/EMG recordings

Surgery was performed 10 days prior baseline recording as described in [64]. 4 days of EEG/EMG signals were annotated on 4s consecutive epochs based on EEG/EMG pattern. Manual annotation was performed on the 3^rd^ day of recording, days 1-2-4 were annotated using a semiautomated scoring system [12, 28].

### Tissue collection

Mice were anesthetized with isoflurane prior to decapitation. Cortex and liver were rapidly dissected, and flash frozen in liquid nitrogen. Time schedule of tissue sampling was described [14].

### RNA-sequencing

Frozen cortex samples were processed as described in [14]. Liver samples were stored at -140°C and prepared as follows: total RNA was extracted using miRNeasy kit (Qiagen; Hilden, Germany). Libraries were prepared using 10 ng/μl with Truseq Stranded RNA. Sequencing was performed on the Illumina HiSeq 4000 SR sequencer with more the 24 million reads per samples.

### Gene quantification from RNA-seq

Gene quantification was performed as follow for both cortex and liver samples: Illumina reads were filtered using fastp [65] to keep high quality reads and remove adapter sequences. Reads were aligned on the mouse reference genome mm10 (GRCm38) using STAR v2.7.0e [66] with default parameters. Read counts was done by STAR using “--quantMode GeneCounts”, taking only reverse strand mapped reads. Genes with low counts (mean counts overall samples < 10) were filtered and normalization was performed with edgeR [67]. Gene expression from the liver was put on Gene Expression Omnibus (GEO) to complete our previous dataset from the cortex. Batch effects were removed using Combat [68] prior fitting using our model.

### Human datasets

Human transcriptome datasets are available on GEO: Forced Desynchrony (GSE48113) and Constant Routine (GSE39445). Experimental details performed are available in the following publications [23, 27]. The following methods are a summary.

### Participants to the Forced Desynchrony

Transcriptome data was obtained from 22 participants (mean ± SD of age, 26.3 ± 3.4 y; 11 males and 11 female). All participants were white, in good health, without reported sleep problems (Pittsburgh Sleep Quality Index ≤5), and homozygous for the PER3 VNTR polymorphism (rs57875989), with equal numbers of ^*4/4*^ and ^*5/5*^ carriers (11 each).

### Forced Desynchrony (FD) protocol

Participants underwent a first 8h baseline sleep schedule at habitual bedtime followed by a 28h sleep-wake cycle. Dark-dim light (<5 lux) cycle and meals also followed a 28h cycle. Plasma melatonin levels were measured as described in [69] to assess circadian period in-vivo and schedule sleep to be in-phase with melatonin levels [70].

### Participants in the Constant Routine

Transcriptome data was obtained from 26 participants (mean ± SD of age, 27.5 ± 4.3 y; 14 males and 12 female). Participants were predominantly white (19/26), in good health, without reported sleep disorder (Pittsburgh Sleep Quality Index ≤5) and homozygous for PER3 VNTR polymorphism (rs57875989).

### Constant Routine (CR) protocol

Participants had to stay awake for 39-41h on their bed, in their individual room in a semi-recumbent position under a low light intensity <10 lux. Hourly nutritional drinks were provided instead of meals. Blood samples were collected hourly to assess melatonin levels and every 3h for total RNA extraction. *Polysomnography*

The EEG, EMG, and EOG (electro-oculogram) were recorded on Siesta 802 devices at a 256Hz sampling rate. After signal filtering, sleep stages were assessed according to Rechtschaffen and Kales criteria. Participants’ sleep was aligned using their melatonin phase and mean sleep amount was calculated using NREM sleep (stages 1-4) + REM sleep and considered baseline sleep onset as “ZT0” in figures.

### RNA extraction, microarray hybridization and processing

Whole peripheral blood was collected using PAXgene Blood RNA tubes. cRNA was hybridized on a 4x44K custom oligonucleotide microarray with additional probes for 20 clock/sleep-related genes. QC and processing were performed with R package limma [71]. Probes intensities were corrected for background and Quantile normalized. Outliers detected with arrayQualityMetrics function and PCA were removed (3/714 samples). For both protocols, blood samples time-point were aligned using participant melatonin phase (i.e., defined as “time point” in FD dataset metadata, and “circadian phase” in CR dataset metadata). Probes were corrected for repeated measure on the same participant using a mixed-model with a random participant intercept and fixed effects of sleep condition (in-phase, anti-phased, 6h sleep + CR, 10h sleep + CR) and time points.

### Driven Damped oscillator model

The temporal dynamic of gene and probes expression were modeled according to the following equation describing a driven damped harmonic oscillator:

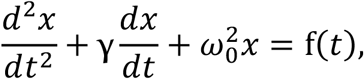

Where *t* is time, *γ* is the linear damping constant and ω_0_ is the natural frequency. Here, we take the the drive f(*t*) as the sum of a drive due to sleep-wake states and a drive due to the master circadian clock in the form of a sinewave. Specifically

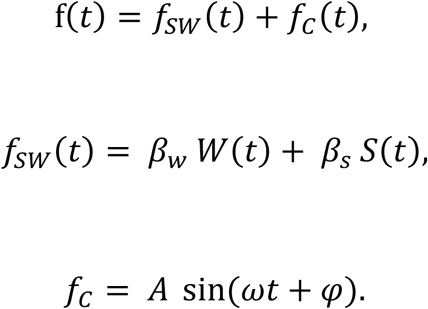

The coefficients *β*_*w*_ and *β*_*s*_ describe the effect of the fraction of sleep and wake per 0.1h bin. *A* and φ are respectively the amplitude and the phase of the circadian drive. The angular velocity ω of the sinewave was set to 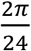, which represents the synchronization of the SCN by the 12:12 light-dark cycle.

### Numerical solution

In order to find optimal parameters for the dynamics of each gene expression we (repeatedly) numerically integrated the driven harmonic oscillator. We first transformed the second order ordinary differential equation (ODE) into two first order ODEs,

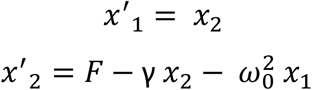

Where *x*_1_*≡ x* and represents normalized mRNA counts and the prime (‘) indicates differentiation with respect to time. We then implemented a 4^th^ order Runge-Kutta (RK4) numerical method to approximate the solution using a fixed time step of 0.1 hour. With a fixed step size of 0.1 hour, RK4 requires values every 0.05 hours. Since values of *fSW*(*t*) were only available every 0.1 hour, we assumed that it took a piecewise constant form.

### Model initial values and optimization procedure

Equilibrium position of the model was set as followed. For each gene or probe, we fitted a cosine to the baseline gene expression (Time 24-48 in mice, FD: in-phase in human) and used the intercept of the model as the default equilibrium position. Initial values of position *x*1(0) and speed *x*2(0) were set at the equilibrium position of the model and at 0, respectively. The baseline sleep-wake cycle (mean baseline sleep in mice, habitual bedtime in human) was repeated for 20 days prior recordings to let the model reach steady state. In humans, an extra free parameter was set for the oscillator equilibrium position in the CR experiment to consider mean difference between FD and CR. This effect could not be corrected in microarray processing directly as no RNA sampling point overlap between experiments, but can be corrected with our model as habitual bedtime sleep are comparable between FD and CR. Optimization was performed using the box-constrained PORT routines method (nlminb) implemented in the optimx/R package. Optimization was done by minimizing the Residual Sum of Square (RSS) between the fit of the model and the expression value of the gene/probe analyzed. A penalization procedure of the RSS was performed to avoid unstable fit in baseline. The maximal and minimal position of the oscillator in the baseline were compared with the position of the oscillator in the 5 days prior baseline (replicated baseline) at the corresponding time. The squared difference was added to the RSS with a weight of 1000. We optimized our model for opposite coefficient sign between sleep and wake and with a minimal 12h period of the natural frequency of our oscillator, to avoid fitting oscillation frequencies too high with respect to gene expression sampling rate. We used multiple starting values for the optimization procedure in an attempt not to reach local optima.

### Model solution

Once optimal parameters were found, we used the analytical solution to decompose the response into the part of the response that was a result of the circadian drive and the part of the response that was a result of the sleep-wake drive, see the Supplementary Material for further details.

### Model Statistics

Goodness of fit was estimated using Kendall’s tau ranked correlation between model fit and expression values. Bayesian Information Criterion (BIC) of the model was calculated from the Negative log likelihood (NLL), assuming that model residuals were independent and followed a Gaussian distribution.

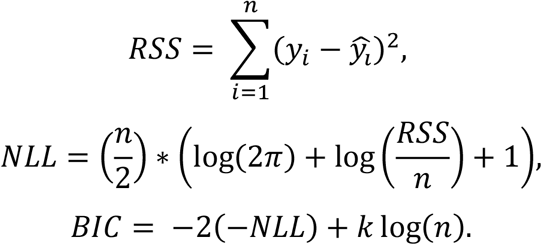

Where n is the number of samples, *y*_*i*_ the gene expression value at time-point i, and k the number of free parameters of the model + 1 (the biased estimator of the error variance 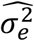. For our model (H_1_): k = 7 for mouse dataset and k=8 for human dataset. For the flat model (H_0_): k=2 for mouse dataset and k=3 for human dataset.

### PCA analysis

PCA analysis in mouse and human and projection of model fitted values were performed using R package FactoMineR [72]. The ellipses were computed using 95% confidence interval of time-points barycentre. In human, missing values were imputed using R package missMDA [73].

### Cortex and liver time delay

To estimate local biological time from clock genes in mouse cortex and liver, we used the R package TimeSignatR (https://github.com/braunr/TimeSignatR) from [60]. Baseline gene expression was used to train the elastic net, penalty parameter alpha and lambda were chosen using a leave-one-out cross validation. Predicted values were obtained from gene expression after sleep deprivation and from model fitted expression.

Using the same strategy, individual gene local biological time was estimated using fitted expression and fitted expression rate in baseline. Expression and Expression rate were fitted to the cartesian coordinate angle of a 24h clock using a bivariate linear model [60]. Genes were filtered for a minimal R^2^ value of the model linear model of 0.6.

## Code Availability

https://github.com/mxjan/SWDMr

## Acknowledgements

We thank Nicolas Guex for his valuable comments on an earlier version of this manuscript. M.J., S.J., and C.N.H. were supported through grants from the Swiss National Science Foundation (31003A_173182 and 310030B_192805 to P.F.). M.J. and P.F. were supported by the University of Lausanne and the Canton de Vaud, Switzerland. D.J.D’s research is supported by the UK Dementia Research Institute which receives its funding from DRI Ltd, funded by the UK Medical Research Council, Alzheimer’s Society and Alzheimer’s Research UK.

## References

1. Zhang, R., et al., A circadian gene expression atlas in mammals: implications for biology and medicine. Proc Natl Acad Sci U S A, 2014. 111(45): p. 16219–24.

2. Mure, L.S., et al., Diurnal transcriptome atlas of a primate across major neural and peripheral tissues. Science, 2018. 359(6381).

3. Talamanca, L., C. Gobet, and F. Naef, Sex-dimorphic and age-dependent organization of 24-hour gene expression rhythms in humans. Science, 2023. 379(6631): p. 478–483.

4. Takahashi, J.S., Transcriptional architecture of the mammalian circadian clock. Nat Rev Genet, 2017. 18(3): p. 164–179.

5. Dibner, C., U. Schibler, and U. Albrecht, The mammalian circadian timing system: organization and coordination of central and peripheral clocks. Annu Rev Physiol, 2010. 72: p. 517–49.

6. Gerber, A., et al., The systemic control of circadian gene expression. Diabetes Obes Metab, 2015. 17 Suppl 1: p. 23–32.

7. Koike, N., et al., Transcriptional architecture and chromatin landscape of the core circadian clock in mammals. Science, 2012. 338(6105): p. 349–54.

8. Kinouchi, K., et al., Circadian rhythms in the tissue-specificity from metabolism to immunity: insights from omics studies. Mol Aspects Med, 2021. 80: p. 100984.

9. Cirelli, C., C.M. Gutierrez, and G. Tononi, Extensive and divergent effects of sleep and wakefulness on brain gene expression. Neuron, 2004. 41(1): p. 35–43.

10. Terao, A., et al., Gene expression in the rat brain during sleep deprivation and recovery sleep: an Affymetrix GeneChip study. Neuroscience, 2006. 137(2): p. 593–605.

11. Hasan, S., et al., How to keep the brain awake? The complex molecular pharmacogenetics of wake promotion. Neuropsychopharmacology, 2009. 34(7): p. 1625–40.

12. Diessler, S., et al., A systems genetics resource and analysis of sleep regulation in the mouse. PLoS Biol, 2018. 16(8): p. e2005750.

13. Mongrain, V., et al., Separating the contribution of glucocorticoids and wakefulness to the molecular and electrophysiological correlates of sleep homeostasis. Sleep, 2010. 33(9): p. 1147–57.

14. Hor, C.N., et al., Sleep-wake-driven and circadian contributions to daily rhythms in gene expression and chromatin accessibility in the murine cortex. Proc Natl Acad Sci U S A, 2019. 116(51): p. 25773–25783.

15. Maret, S., et al., Homer1a is a core brain molecular correlate of sleep loss. Proc Natl Acad Sci U S A, 2007. 104(50): p. 20090–5.

16. Suzuki, A., M. Yanagisawa, and R.W. Greene, Loss of Arc attenuates the behavioral and molecular responses for sleep homeostasis in mice. Proc Natl Acad Sci U S A, 2020. 117(19): p. 10547–10553.

17. Tononi, G. and C. Cirelli, Sleep and synaptic down-selection. Eur J Neurosci, 2020. 51(1): p. 413–421.

18. Diering, G.H., Remembering and forgetting in sleep: Selective synaptic plasticity during sleep driven by scaling factors Homer1a and Arc. Neurobiol Stress, 2023. 22: p. 100512.

19. Franken, P., et al., A non-circadian role for clock-genes in sleep homeostasis: a strain comparison. BMC Neurosci, 2007. 8: p. 87.

20. Wang, H., et al., Computational analysis of gene regulation in animal sleep deprivation. Physiol Genomics, 2010. 42(3): p. 427–36.

21. Franken, P., A role for clock genes in sleep homeostasis. Curr Opin Neurobiol, 2013. 23(5): p. 864–72.

22. Noya, S.B., et al., The forebrain synaptic transcriptome is organized by clocks but its proteome is driven by sleep. Science, 2019. 366(6462).

23. Archer, S.N., et al., Mistimed sleep disrupts circadian regulation of the human transcriptome. Proc Natl Acad Sci U S A, 2014. 111(6): p. E682–91.

24. Maury, E., H.K. Hong, and J. Bass, Circadian disruption in the pathogenesis of metabolic syndrome. Diabetes Metab, 2014. 40(5): p. 338–46.

25. Roenneberg, T. and M. Merrow, The Circadian Clock and Human Health. Curr Biol, 2016. 26(10): p. R432–43.

26. Daan, S., D.G. Beersma, and A.A. Borbely, Timing of human sleep: recovery process gated by a circadian pacemaker. Am J Physiol, 1984. 246(2 Pt 2): p. R161–83.

27. Moller-Levet, C.S., et al., Effects of insufficient sleep on circadian rhythmicity and expression amplitude of the human blood transcriptome. Proc Natl Acad Sci U S A, 2013. 110(12): p. E1132–41.

28. Jan, M., et al., A multi-omics digital research object for the genetics of sleep regulation. Sci Data, 2019. 6(1): p. 258.

29. Archer, S.N., et al., Mistimed sleep and waking activity in humans disrupts glucocorticoid signalling transcripts and SP1, but not plasma cortisol rhythms. Front Physiol, 2022. 13: p. 946444.

30. Masubuchi, S., et al., Clock genes outside the suprachiasmatic nucleus involved in manifestation of locomotor activity rhythm in rats. Eur J Neurosci, 2000. 12(12): p. 4206–14.

31. Abe, H., et al., Behavioural rhythm splitting in the CS mouse is related to clock gene expression outside the suprachiasmatic nucleus. Eur J Neurosci, 2001. 14(7): p. 1121–8.

32. Wakamatsu, H., et al., Restricted-feeding-induced anticipatory activity rhythm is associated with a phase-shift of the expression of mPer1 and mPer2 mRNA in the cerebral cortex and hippocampus but not in the suprachiasmatic nucleus of mice. Eur J Neurosci, 2001. 13(6): p. 1190–6.

33. Curie, T., et al., In Vivo Imaging of the Central and Peripheral Effects of Sleep Deprivation and Suprachiasmatic Nuclei Lesion on PERIOD-2 Protein in Mice. Sleep, 2015. 38(9): p. 1381–94.

34. Deboer, T., et al., Sleep states alter activity of suprachiasmatic nucleus neurons. Nat Neurosci, 2003. 6(10): p. 1086–90.

35. Challet, E., et al., Sleep deprivation decreases phase-shift responses of circadian rhythms to light in the mouse: role of serotonergic and metabolic signals. Brain Res, 2001. 909(1-2): p. 81–91.

36. Antle, M.C. and R.E. Mistlberger, Circadian clock resetting by sleep deprivation without exercise in the Syrian hamster. J Neurosci, 2000. 20(24): p. 9326–32.

37. Gekakis, N., et al., Role of the CLOCK protein in the mammalian circadian mechanism. Science, 1998. 280(5369): p. 1564–9.

38. Zhang, Y., et al., Targeted deletion of thioesterase superfamily member 1 promotes energy expenditure and protects against obesity and insulin resistance. Proc Natl Acad Sci U S A, 2012. 109(14): p. 5417–22.

39. Yin, L. and M.A. Lazar, The orphan nuclear receptor Rev-erbalpha recruits the N-CoR/histone deacetylase 3 corepressor to regulate the circadian Bmal1 gene. Mol Endocrinol, 2005. 19(6): p. 1452–9.

40. Alenghat, T., et al., Nuclear receptor corepressor and histone deacetylase 3 govern circadian metabolic physiology. Nature, 2008. 456(7224): p. 997–1000.

41. Aninye, I.O., et al., Circadian regulation of Tshb gene expression by Rev-Erbalpha (NR1D1) and nuclear corepressor 1 (NCOR1). J Biol Chem, 2014. 289(24): p. 17070–7.

42. Mistlberger, R.E., et al., Recovery sleep following sleep deprivation in intact and suprachiasmatic nuclei-lesioned rats. Sleep, 1983. 6(3): p. 217–33.

43. Herzog, E.D., et al., Regulating the Suprachiasmatic Nucleus (SCN) Circadian Clockwork: Interplay between Cell-Autonomous and Circuit-Level Mechanisms. Cold Spring Harb Perspect Biol, 2017. 9(1).

44. Gerstner, J.R., et al., Removal of unwanted variation reveals novel patterns of gene expression linked to sleep homeostasis in murine cortex. BMC Genomics, 2016. 17(Suppl 8): p. 727.

45. Curie, T., et al., Homeostatic and circadian contribution to EEG and molecular state variables of sleep regulation. Sleep, 2013. 36(3): p. 311–23.

46. Hoekstra, M.M., et al., The sleep-wake distribution contributes to the peripheral rhythms in PERIOD-2. Elife, 2021. 10.

47. Sneppen, K., S. Krishna, and S. Semsey, Simplified models of biological networks. Annu Rev Biophys, 2010. 39: p. 43–59.

48. De Los Santos, H., K.P. Bennett, and J.M. Hurley, MOSAIC: a joint modeling methodology for combined circadian and non-circadian analysis of multi-omics data. Bioinformatics, 2021. 37(6): p. 767–774.

49. Spiess, A.N. and N. Neumeyer, An evaluation of R2 as an inadequate measure for nonlinear models in pharmacological and biochemical research: a Monte Carlo approach. BMC Pharmacol, 2010. 10: p. 6.

50. Miller, B.H., et al., Circadian and CLOCK-controlled regulation of the mouse transcriptome and cell proliferation. Proc Natl Acad Sci U S A, 2007. 104(9): p. 3342–7.

51. Seon Je-young, K.W.-s., Kim Jong-woo, Lim Seong-bin, Oh Mi-ae, Changes in Human Gene Expression After Sleep Deprivation. Korean J Biol Psychiatry, 2022. 29(1): p. p9–14.

52. Wittenbrink, N., et al., High-accuracy determination of internal circadian time from a single blood sample. J Clin Invest, 2018. 128(9): p. 3826–3839.

53. Moller-Levet, C.S., et al., Diurnal and circadian rhythmicity of the human blood transcriptome overlaps with organ- and tissue-specific expression of a non-human primate. BMC Biol, 2022. 20(1): p. 63.

54. Laing, E.E., et al., Identifying and validating blood mRNA biomarkers for acute and chronic insufficient sleep in humans: a machine learning approach. Sleep, 2019. 42(1).

55. Koh, K., et al., Identification of SLEEPLESS, a sleep-promoting factor. Science, 2008. 321(5887): p. 372–6.

56. Chua, E.C., et al., Changes in Plasma Lipids during Exposure to Total Sleep Deprivation. Sleep, 2015. 38(11): p. 1683–91.

57. Ashlin, T.G., et al., Pitpnc1a Regulates Zebrafish Sleep and Wake Behavior through Modulation of Insulin-like Growth Factor Signaling. Cell Rep, 2018. 24(6): p. 1389–1396.

58. Bolsius, Y.G., et al., The role of clock genes in sleep, stress and memory. Biochem Pharmacol, 2021. 191: p. 114493.

59. Reinke, H. and G. Asher, Crosstalk between metabolism and circadian clocks. Nat Rev Mol Cell Biol, 2019. 20(4): p. 227–241.

60. Braun, R., et al., Universal method for robust detection of circadian state from gene expression. Proc Natl Acad Sci U S A, 2018. 115(39): p. E9247–E9256.

61. Dijk, D.J. and J.F. Duffy, Novel Approaches for Assessing Circadian Rhythmicity in Humans: A Review. J Biol Rhythms, 2020. 35(5): p. 421–438.

62. Borbely, A.A., A two process model of sleep regulation. Hum Neurobiol, 1982. 1(3): p. 195–204.

63. Hafycz, J.M., E. Strus, and N. Naidoo, Reducing ER stress with chaperone therapy reverses sleep fragmentation and cognitive decline in aged mice. Aging Cell, 2022. 21(6): p. e13598.

64. Mang, G.M. and P. Franken, Sleep and EEG Phenotyping in Mice. Curr Protoc Mouse Biol, 2012. 2(1): p. 55–74.

65. Chen, S., et al., fastp: an ultra-fast all-in-one FASTQ preprocessor. Bioinformatics, 2018. 34(17): p. i884–i890.

66. Dobin, A., et al., STAR: ultrafast universal RNA-seq aligner. Bioinformatics, 2013. 29(1): p. 15–21.

67. Robinson, M.D., D.J. McCarthy, and G.K. Smyth, edgeR: a Bioconductor package for differential expression analysis of digital gene expression data. Bioinformatics, 2010. 26(1): p. 139–40.

68. Johnson, W.E., C. Li, and A. Rabinovic, Adjusting batch effects in microarray expression data using empirical Bayes methods. Biostatistics, 2007. 8(1): p. 118–27.

69. Hasan, S., et al., Assessment of circadian rhythms in humans: comparison of real-time fibroblast reporter imaging with plasma melatonin. FASEB J, 2012. 26(6): p. 2414–23.

70. Lazar, A.S., et al., Circadian period and the timing of melatonin onset in men and women: predictors of sleep during the weekend and in the laboratory. J Sleep Res, 2013. 22(2): p. 155–9.

71. Ritchie, M.E., et al., limma powers differential expression analyses for RNA-sequencing and microarray studies. Nucleic Acids Res, 2015. 43(7): p. e47.

72. Lê, S., J. Josse, and F. Husson, FactoMineR: An R Package for Multivariate Analysis. Journal of Statistical Software, 2008. 25(1): p. 1–18.

73. Josse, J. and F. Husson, missMDA: A Package for Handling Missing Values in Multivariate Data Analysis. Journal of Statistical Software, 2016. 70(1): p. 1–31.

