## Supplementary Material for "Model integration of circadian and sleep-wake driven contributions to rhythmic gene expression reveals novel regulatory principles"

Maxime Jan[1,2], Sonia Jimenez[1], Charlotte N Hor[1], Derk-Jan Dijk[3,4], Anne C Skeldon[4,5], and Paul Franken[1]

[1]Center of Integrative Genomics, University of Lausanne, Switzerland
[2]Bioinformatics Competence Center, University of Lausanne, Switzerland
[3]Surrey Sleep Research Centre, University of Surrey, Guildford, UK
[4]UK Dementia Research Institute, Care Research and Technology Centre at Imperial College, London and the University of Surrey, Guildford, UK
[5]Department of Mathematics, University of Surrey, Guildford, UK

August 9, 2023

#### Derivation of the driven damped oscillator model

Our goal was to develop a simple model which had the ability to exhibit both exponential and oscillatory behaviour. The driven damped harmonic oscillator model is one such model. Here we add additional details to further explain how the driven damped harmonic oscillator equation can result from a simple model of the interaction of a gene with its environment and external driving factors. In the manuscript we state

$$\begin{aligned} \frac{dX}{dt} &= \alpha Y - \gamma X, \\ \frac{dY}{dt} &= -\beta X + F(t), \end{aligned} \tag{S1}$$

where $X(t)$ is the level of mRNA of a gene, $Y(t)$ is a combination of intra-tissue factors which we term 'tissue environment', and $F(t)$ captures external driving factors including sleep and wake and circadian rhythmicity. The constants $\alpha, \beta, \gamma$ describe the effect of the environment on the gene, the effect of the gene on the environment and the gene degradation rate respectively.

We let $X(t) = X_b + x(t)$, $Y(t) = Y_b + y(t)$, $F(t) = F_b + f(t)$, where $x(t), y(t), f(t)$ model the deviation from a stationary baseline. Substituting $X(t)$, $Y(t)$ and $F(t)$ into equations (S1) gives

$$\begin{aligned} \frac{dx}{dt} &= \alpha y - \gamma x + (\alpha Y_b - \gamma X_b), \\ \frac{dy}{dt} &= -\beta x + f(t) + (-\beta X_b + F_b). \end{aligned} \tag{S2}$$

Since the stationary baseline values satisfy equations (S1) with $dX/dt = dY/dt = 0$, the terms in the brackets on the right hand side of equations (S2) are zero, leaving

$$\frac{dx}{dt} = \alpha y - \gamma x, \tag{S3}$$

$$\frac{dy}{dt} = -\beta x + f(t). \tag{S4}$$

We note that while we would expect $X(t) > 0$, $x(t)$ may be positive or negative.

Now, differentiating equation (S3) with respect to time gives

$$\frac{d^2x}{dt^2} = \alpha\frac{dy}{dt} - \gamma\frac{dx}{dt},$$

Substituting for $dy/dt$ from equation (S4) and re-arranging gives

$$\frac{d^2x}{dt^2} + \gamma\frac{dx}{dt} + \alpha\beta\, x = \alpha f(t),$$

i.e. the equation of a damped oscillator with natural frequency $\omega_0 = \sqrt{\alpha\beta}$ driven by $\alpha f(t)$.

### Solutions of the driven damped oscillator model

Consider the driven damped harmonic oscillator

$$\frac{d^2x}{dt^2} + \gamma\frac{dx}{dt} + \omega_0^2 x = C + F_1\cos\omega t + F_2\sin\omega t, \tag{S5}$$

with initial conditions

$$x(0) = \alpha, \qquad \left.\frac{dx}{dt}\right|_{t=0} = \beta,$$

where we note that the right hand side of equation (S5) could alternatively be written as

$$C + A\sin(\omega t + \phi),$$

where $F_1 \equiv A\sin\phi, F_2 \equiv A\cos\phi$. Solutions to equation (S5) may be found using standard methods of calculus and consist of a linear combination of the 'complementary function' $x_{CF}(t)$, which is the solution to the homogeneous equation, namely

$$\frac{d^2x}{dt^2} + \gamma\frac{dx}{dt} + \omega_0^2 x = 0 \tag{S6}$$

and the 'particular integral' which is one solution to the inhomogeneous equation (S5). Here we separate the particular integral into two parts, $x_h$ and $x_c(t)$ where $x_h$ is the consequence of a constant driving term $C$ and $x_c(t)$ is the consequence of the oscillatory driving term $F_1\cos\omega t + F_2\sin\omega t$. Hence the solution to equation (S5) takes the form

$$x(t) = x_{CF}(t) + x_h + x_c(t). \tag{S7}$$

The particular solution for the constant driving term is given by

$$x_h = \frac{C}{\omega_0^2}, \tag{S8}$$

and that for the two oscillatory terms as

$$x_c = D\cos\omega t + E\sin\omega t \equiv F\sin(\omega t + \tilde{\phi}), \tag{S9}$$

where

$$D = \frac{\left(\omega_0^2 - \omega^2\right)F_2 - \gamma\omega F_1}{\gamma^2\omega^2 + \left(\omega_0^2 - \omega^2\right)^2} \qquad \text{and} \qquad E = \frac{\gamma\omega F_2 + \left(\omega_0^2 - \omega^2\right)F_1}{\gamma^2\omega^2 + \left(\omega_0^2 - \omega^2\right)^2}, \tag{S10}$$

and $F = \sqrt{D^2 + E^2}, \tan\tilde{\phi} = D/E$.

The solution to the homogeneous equation, $x_{CF}$, separates into three types depending on the sign of $\gamma^2 - 4\omega_0^2$. Specifically, if $\gamma^2 - 4\omega_0^2 > 0$, solutions are 'overdamped' and

$$x_{CF} = A_{\mathrm{o}}\mathrm{e}^{a_1 t} + B_{\mathrm{o}}\mathrm{e}^{a_1 t}, \tag{S11}$$

where

$$a_{1,2} = \frac{-\gamma \pm \sqrt{\gamma^2 - 4\omega_0^2}}{2}$$

are real and negative.

If $\gamma^2 = 4\omega_0^2$, solutions are 'critically damped',

$$x_{CF} = (A_{\mathrm{c}}t + B_{\mathrm{c}})\,\mathrm{e}^{-\gamma t/2}. \tag{S12}$$

Finally, if $\gamma^2 - 4\omega_0^2 < 0$, solutions are 'underdamped',

$$x_{CF} = (A_{\mathrm{u}}\cos\omega_1 t + B_{\mathrm{u}}\sin\omega_1 t)\,\mathrm{e}^{-\gamma t/2}, \tag{S13}$$

where $\omega_1^2 = \omega_0^2 - \gamma^2/4$. We note that it is a matter of convention that the solutions for the overdamped and underdamped cases are written as exponentials / trigonometric functions respectively. Specifically, solutions for the underdamped case may be written in the form of equation (S12) but with $a_{1,2}$ complex instead of real. Similarly, the overdamped case may be written in the form of equation (S12) but the frequency $\omega_1$ will be imaginary instead of real. The fact that the two forms are equivalent is relevant for our separation of circadian and sleep/wake effects described below, as it means that a single formulation covers both cases.

Values for the constants $A_{\mathrm{o}}, B_{\mathrm{o}}$ or $A_{\mathrm{c}}, B_{\mathrm{c}}$ or $A_{\mathrm{u}}, B_{\mathrm{u}}$ may be found using the initial conditions. For example,

$$\begin{aligned} A_{\mathrm{u}} &= \alpha - \frac{C}{\omega_0^2} - D, \\ B_{\mathrm{u}} &= \frac{\beta + \frac{\gamma}{2}\left(\alpha - \frac{C}{\omega_0^2} - D\right) - \omega E}{\omega_1}. \end{aligned}$$

Hence, the solution is completely determined by the seven constants in equation (S5) namely the damping parameters $\gamma$, the natural frequency $\omega_0$, the size of the constant driving term $C$, the two parameters specifying the sinusoidal driving term, either the amplitude and phase ($A$ and $\phi$) or equivalently $F_1$ and $F_2$ and the two values specifying the initial conditions $\alpha$ and $\beta$. The solution consists of three parts, where

$x_h$ is a constant, $x_c(t)$ is oscillatory with a fixed amplitude and phase. In the underdamped case $x_{CF}(t)$, is a damped oscillatory term (see equation (S13)) so

$$x(t) = x_{CF}(t) + x_h + x_c(t) = (A_\mathrm{u}\cos\omega_1 t + B_\mathrm{u}\sin\omega_1 t)\,\mathrm{e}^{-\gamma t/2} + \frac{C}{\omega_0^2} + D\cos\omega t + E\sin\omega t. \tag{S14}$$

In the overdamped case $x_{CF}(t)$ is non-oscillatory (see equation (S12)), and instead the general solution takes the form

$$x(t) = A_\mathrm{o}\mathrm{e}^{a_1 t} + B_\mathrm{o}\mathrm{e}^{a_1 t} + \frac{C}{\omega_0^2} + D\cos\omega t + E\sin\omega t. \tag{S15}$$

Finally, in the critically damped case the general solution takes the form

$$x(t) = (A_\mathrm{c} t + B_\mathrm{c})\,\mathrm{e}^{-\gamma t/2} + \frac{C}{\omega_0^2} + D\cos\omega t + E\sin\omega t. \tag{S16}$$

### Process-S-like dynamics in the driven damped oscillator

Process-S-like sleep-wake driven processes are usually described as exponential functions of the form

$$S(t) = \tilde{C} - \left(\tilde{C} - S_0\right)\mathrm{e}^{-kt},$$

where $S(t)$ is homeostatic sleep pressure, $\tilde{C}$ is the asymptote i.e. $S(t) \to \tilde{C}$ as $t \to \infty$, $k$ is the decay rate and $S(t) = S_0$ at $t = 0$. In the absence of a sinusoidal driving term, the solution for the critically damped driven oscillator, given in equation (S16), reduces to $S(t)$ for the initial conditions $x(0) = S_0$ and $x'(0) = k(\tilde{C} - S_0)$.

### Piecewise constant driving

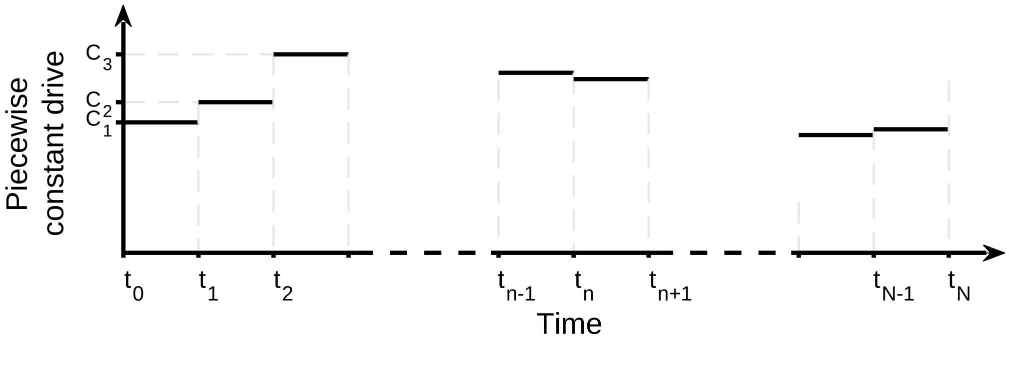


Figure S1: Piecewise constant drive as a result of sleep and wake.

Sleep is assumed to result in a piecewise constant driving term, such that for $t \in (t_{n-1}, t_n)$, where $n$ is the interval number $n = 1, 2\ldots$ it takes the value $C_n$ where the magnitude of the $C_n$ is directly proportional to the fraction of time asleep. Taking the initial conditions for the $n^{th}$ interval as

$$x(t_{n-1}) = \alpha_{n-1}, \qquad \left.\frac{dx}{dt}\right|_{t=t_{n-1}} = \beta_{n-1}.$$

and using equation (S5), then for the underdamped case the solution for $x(t)$ in the $n^{th}$ interval, $x_n(t)$, has the general form

$$x_n(t) = (A_n \cos\omega_1 t + B_n \sin\omega_1 t)\,\mathrm{e}^{-\gamma t/2} + \frac{C_n}{\omega_0^2} + D\cos\omega t + E\sin\omega t, \tag{S17}$$

where $D$ and $E$ are

$$D = \frac{\left(\omega_0^2 - \omega^2\right) F_2 - \gamma\omega F_1}{\gamma^2\omega^2 + \left(\omega_0^2 - \omega^2\right)^2} \qquad \text{and} \qquad E = \frac{\gamma\omega F_2 + \left(\omega_0^2 - \omega^2\right) F_1}{\gamma^2\omega^2 + \left(\omega_0^2 - \omega^2\right)^2}, \tag{S18}$$

as in (S10), and $A_n$ and $B_n$ satisfy the linear simultaneous equations

$$\begin{aligned}
\alpha_{n-1} &= (A_n \cos\omega_1 t_{n-1} + B_n \sin\omega_1 t_{n-1})\,\mathrm{e}^{-\gamma t_{n-1}/2} + \frac{C_n}{\omega_0^2} + D\cos\omega t_{n-1} + E\sin\omega t_{n-1}, \\
\beta_{n-1} &= -\frac{\gamma}{2}\,(A_n \cos\omega_1 t_{n-1} + B_n \sin\omega_1 t_{n-1})\,\mathrm{e}^{-\gamma t_{n-1}/2} \\
&\quad + (-\omega_1 A_n \sin\omega_1 t_{n-1} + \omega_1 B_n \cos\omega_1 t_{n-1})\,\mathrm{e}^{-\gamma t_{n-1}/2} - \omega D \sin\omega t_{n-1} + E\omega\cos\omega t_{n-1}.
\end{aligned} \tag{S19}$$

Solving equations (S19) gives

$$\begin{aligned}
A_n &= \frac{\mathrm{e}^{\gamma t_{n-1}/2}}{\omega_1}\left\{\left(-\frac{\gamma}{2}\sin\omega_1 t_{n-1} + \omega_1\cos\omega_1 t_{n-1}\right)\left(\alpha_{n-1} - \frac{C_n}{\omega_0^2}\right) - \sin\omega_1 t_{n-1}\,\beta_{n-1}\right. \\
&\quad + \left(\frac{\gamma}{2}\sin\omega_1 t_{n-1} - \omega_1\cos\omega_1 t_{n-1}\right)(D\cos\omega t_{n-1} + E\sin\omega t_{n-1}) \\
&\quad\left. - \,\omega\sin\omega_1 t_{n-1}\,(D\sin\omega t_{n-1} - E\cos\omega t_{n-1})\right\} \\
B_n &= \frac{\mathrm{e}^{\gamma t_{n-1}/2}}{\omega_1}\left\{\left(\frac{\gamma}{2}\cos\omega_1 t_{n-1} + \omega_1\sin\omega_1 t_{n-1}\right)\left(\alpha_{n-1} - \frac{C_n}{\omega_0^2}\right) + \cos\omega_1 t_{n-1}\,\beta_{n-1}\right. \\
&\quad - \left(\frac{\gamma}{2}\cos\omega_1 t_{n-1} + \omega_1\sin\omega_1 t_{n-1}\right)(D\cos\omega t_{n-1} + E\sin\omega t_{n-1}) \\
&\quad\left. + \,\omega\cos\omega_1 t_{n-1}\,(D\cos\omega t_{n-1} - E\sin\omega t_{n-1})\right\}
\end{aligned} \tag{S20}$$

Hence, given $\omega_0, \gamma$, a constant $C$ which relates faction of time asleep to the piecewise constant drive, i.e. $C_n = C\times$ fraction of time asleep, the amplitude $A$, phase $\phi$ and angular frequency $\omega$ (here, $2\pi/24$ radians / hour) of the oscillatory driving term the solution is calculated as follows.

- From $\omega_0$ and $\gamma$ calculate $\omega_1 = \surd(\omega_0^2 - \gamma^2)$.
- From $A$ and $\phi$ calculate the values of $F_1$ and $F_2$ since $F_1 = A\sin\phi, F_2 = A\cos\phi$.
- Calculate $D$ and $E$ from equation (S18).
- Work iteratively through each time interval, starting at $t_0$. For each interval the solution $x_n(t)$ is derived by evaluating $A_n$, $B_n$ using equations (S20). Once $x_n(t)$ is found, the starting conditions $\alpha_n$ and $\beta_n$ for the next interval may be found by evaluating $x_n(t_n)$ and $x'_n(t_n)$ where $x'_n(t)$ is the derivative of $x_n(t)$ with respect to time.

In order to start this iterative process, values for $\alpha_0$ and $\beta_0$ are required. In order to ensure that the results were insensitive to the choice of $\alpha_0$ and $\beta_0$, the sleep and circadian drives were prepended by 20 replicates of the baseline day.

The five required constants $\omega_0, \gamma, C, A$ and $\phi$ could be found by fitting the analytical solution to the gene expression data. In practice, these constants were evaluated instead by numerically integrating the oscillator equations using a fourth order Runge-Kutta method with a fixed step size. For a given set of constants, the numerical solution and the analytical solution matched to a high degree of accuracy (typical error less than $10^{-7}$, close to the precision of the variables), suggesting that any numerical errors are negligible. We note that the fitting was done with a piecewise constant sleep-wake drive, which in the n$^{th}$ interval is given by $\beta_W W_n(t) + \beta_S S_n(t)$, rather than using a sleep drive, as described in the analytical solution above. However, it is straightforward to transform between the two alternatives since in each 0.1 h interval the mouse is either asleep or awake, so $W_n(t) + S_n(t) = 0.1$. Hence the sleep-wake drive may alternatively be formulated as a constant plus sleep drive, i.e. $0.1\beta_W + (\beta_W - \beta_S)S_n(t)$.

The numerical solution yields a single fitted time series for each gene. The analytical solution then enables that single time trace to be separated into components driven by the different driving terms equivalent to the $x_{CF}(t)$, $x_h$ and $x_c(t)$ in equation (S14), see Fig. S2 for one example. Specifically, the term $x_c(t)$ is the response to the oscillatory drive. The term $x_h$ is a response to the (piecewise) constant drive. The $x_{CF}(t)$ is normally termed as the 'transient' since $x_{CF}(t) \to 0$ as $t \to \infty$. Here, we find that $x_{CF}(t)$ responds to the short timescale changes in fraction of time asleep so is not negligible. Consequently, in decomposing the fitted timeseries into circadian and homeostatic contributions, we consider that circadian contributions are given by $x_c(t)$ and homeostatic contributions are given by $x_h + x_{CF}(t)$.

The method used to calculate $\omega_1, D, E, \alpha_n, \beta_n, A_n$ and $B_n$ was described above for the underdamped case and coded in MATLAB. The same piece of code also worked for the overdamped case since, as discussed above, it is a matter of convention rather than a fundamental difference in the mathematical formula that distinguishes the two cases.

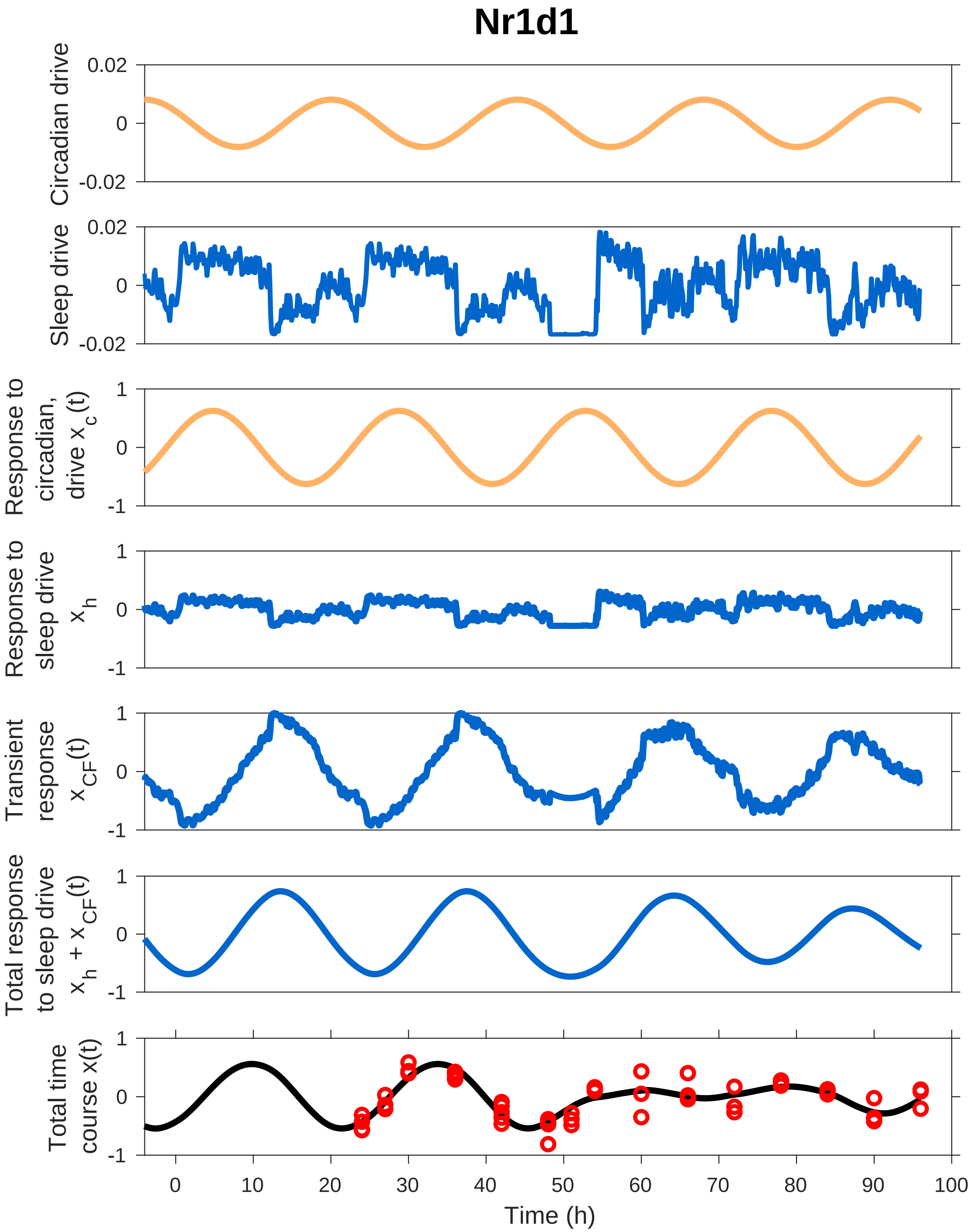


Figure S2: Circadian and sleep driving terms and consequent solution components for one example (Nr1d1). The fitted gene expression data are shown in red.
