## Supplementary material for "Model integration of circadian and sleep-wake driven contributions to rhythmic gene expression reveals novel regulatory principles": Fig. S5

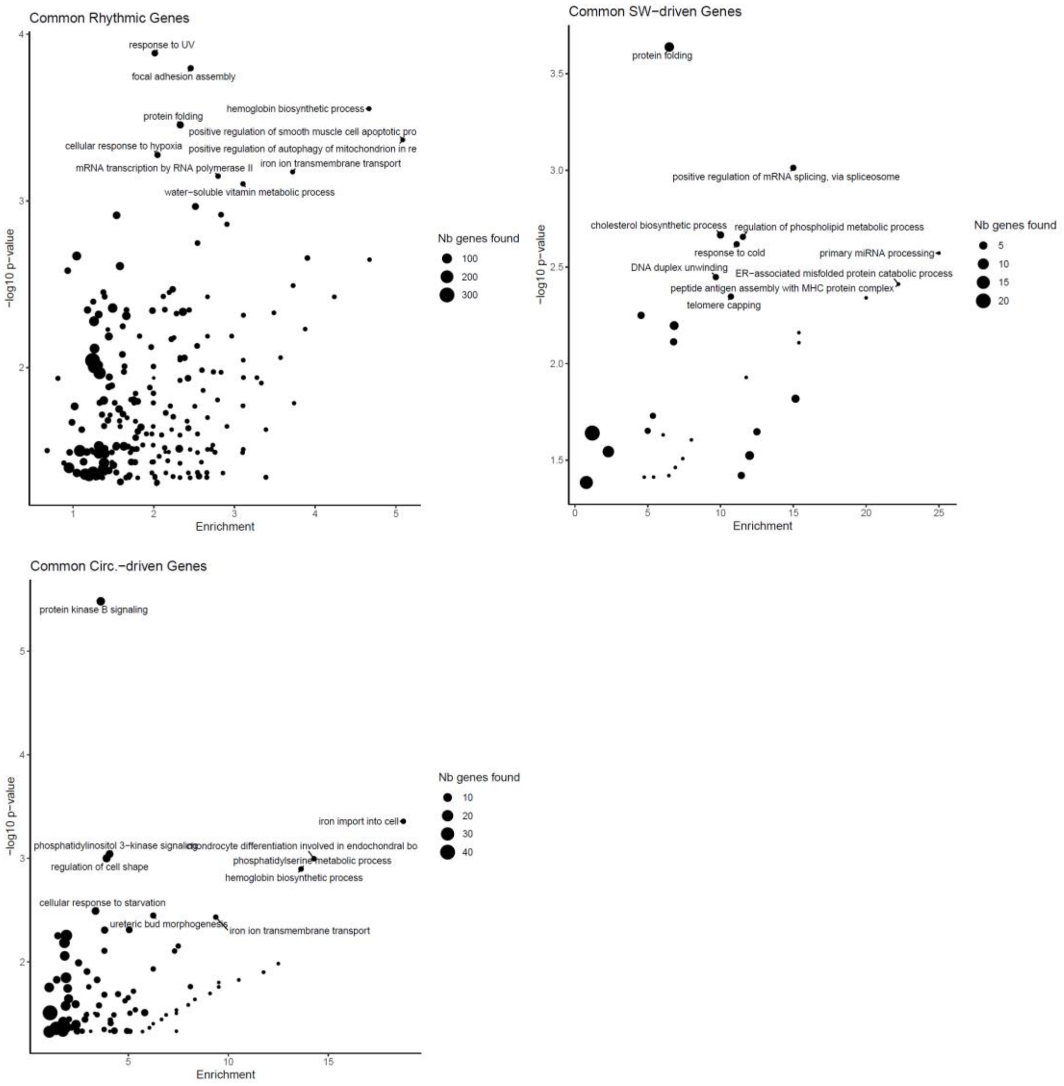


**Figure S5: GO enrichment for common sleep-wake or circadian driven genes.** GO term enrichment for common rhythmic genes ($\Delta BIC_{10} > 2$; n=1468; top left), for common sleep-wake driven genes (*SWrc* > 0.5; n=109; top right), and for common circadian driven (*SWrc* > 0.5; n=215; bottom) genes in mouse cortex, liver, and human blood (intersection of tissues; see **Fig. 6B**). As background the intersection of genes expressed in cortex, liver, and blood was used in the GO-enrichment analysis.
