## Supplementary material for "Model integration of circadian and sleep-wake driven contributions to rhythmic gene expression reveals novel regulatory principles": Fig. S6

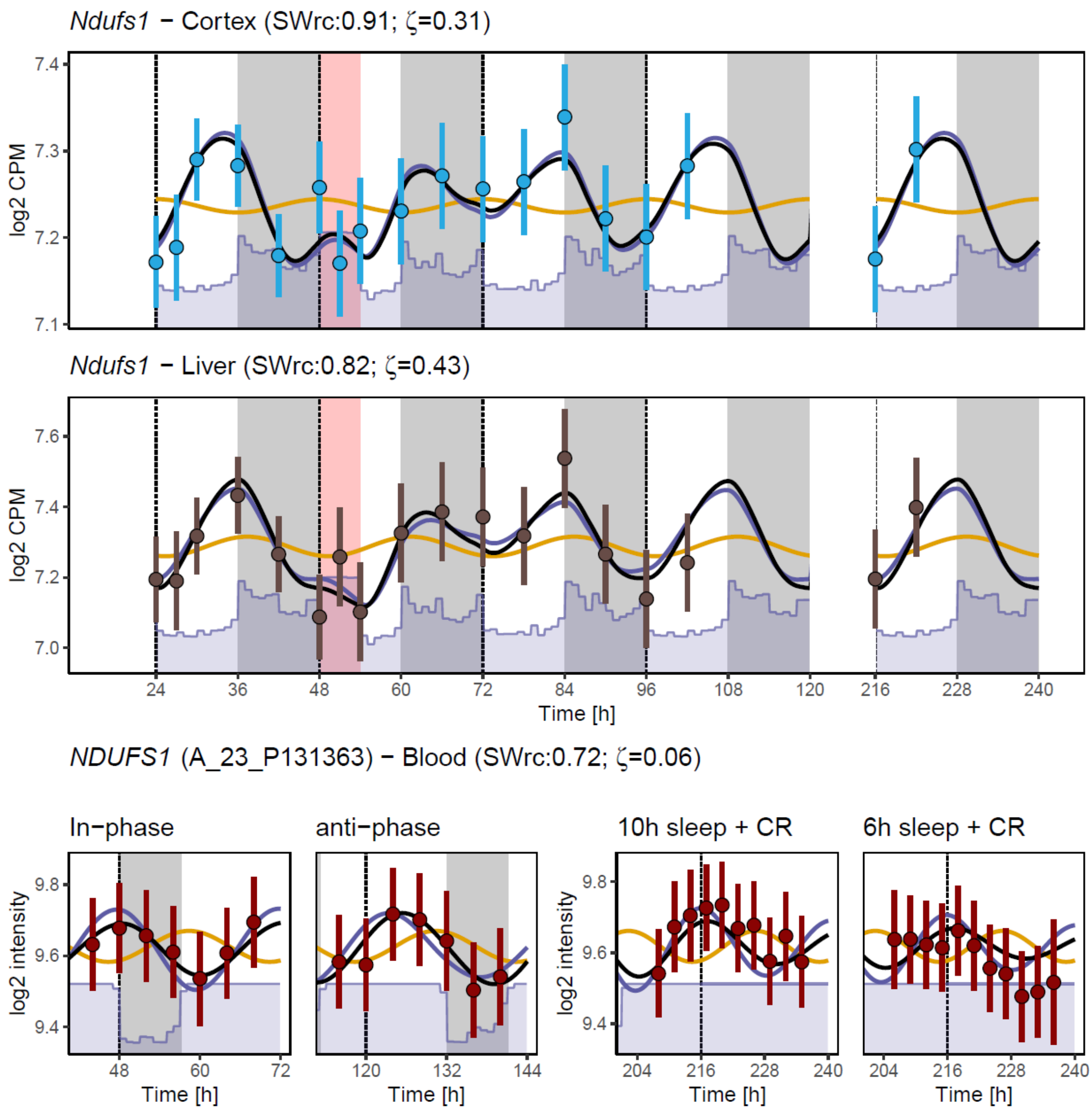


**Figure S6: Common sleep-wake driven genes in liver, cortex, and blood:** Among the 109 genes labelled as sleep-wake driven (*SWrc* > 0.5) in all 3 tissues (cortex=blue, liver = brown, blood = red) *Ndufs1* had the highest average *SWrc* (0.82). Fitted expression as black line, circadian response as orange line, and sleep-wake response as blue line.
