## Supplementary material for "Model integration of circadian and sleep-wake driven contributions to rhythmic gene expression reveals novel regulatory principles": Fig. S1

### Supplemental Figures

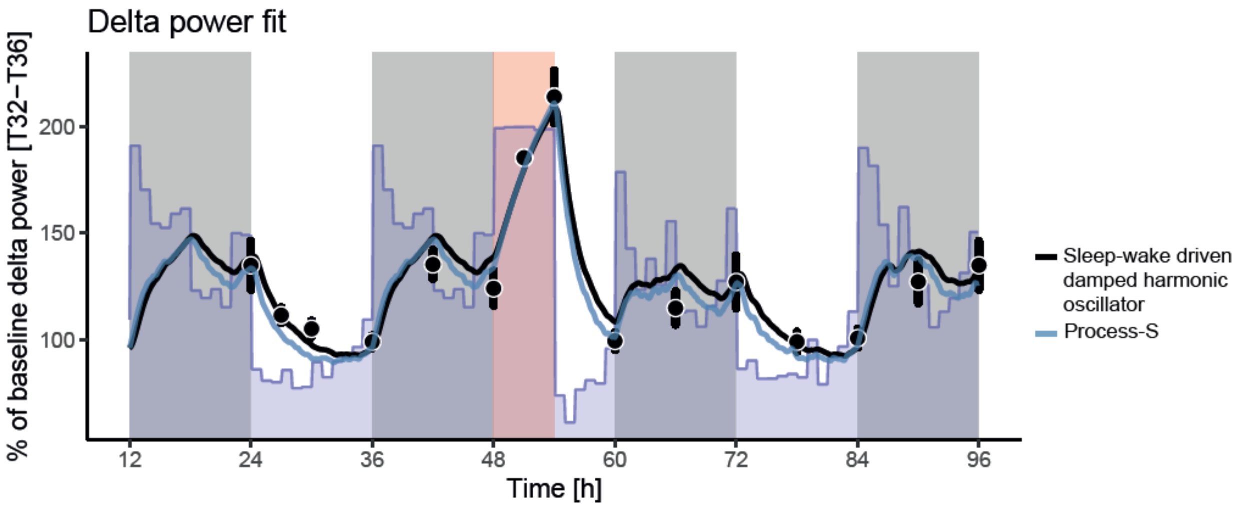


**Figure S1: A sleep-wake driven damped harmonic oscillator can accurately fit the dynamics EEG delta power [1-4Hz] in mice.** Black symbols: observed delta-power in NREM sleep expressed as % of lowest values reached in baseline. Blue line: simulation of EEG delta power using the 'classical' model using exponential saturating functions increasing during wakefulness and REM sleep and decreasing during NREM sleep using previously obtained parameters "Process S" in (Franken, Dudley et al. 2006). Black line: fitted overdamped harmonic oscillator with time spent in wake and REM sleep as positive forces accelerating and NREM sleep as negative force decelerating delta power. For the oscillator model, fitting was done on log transformed values of delta-power % and then transformed back to % values for plot. Blue area hourly values of Wake + REM sleep.
