## Supplementary material for "Model integration of circadian and sleep-wake driven contributions to rhythmic gene expression reveals novel regulatory principles": Fig. S2

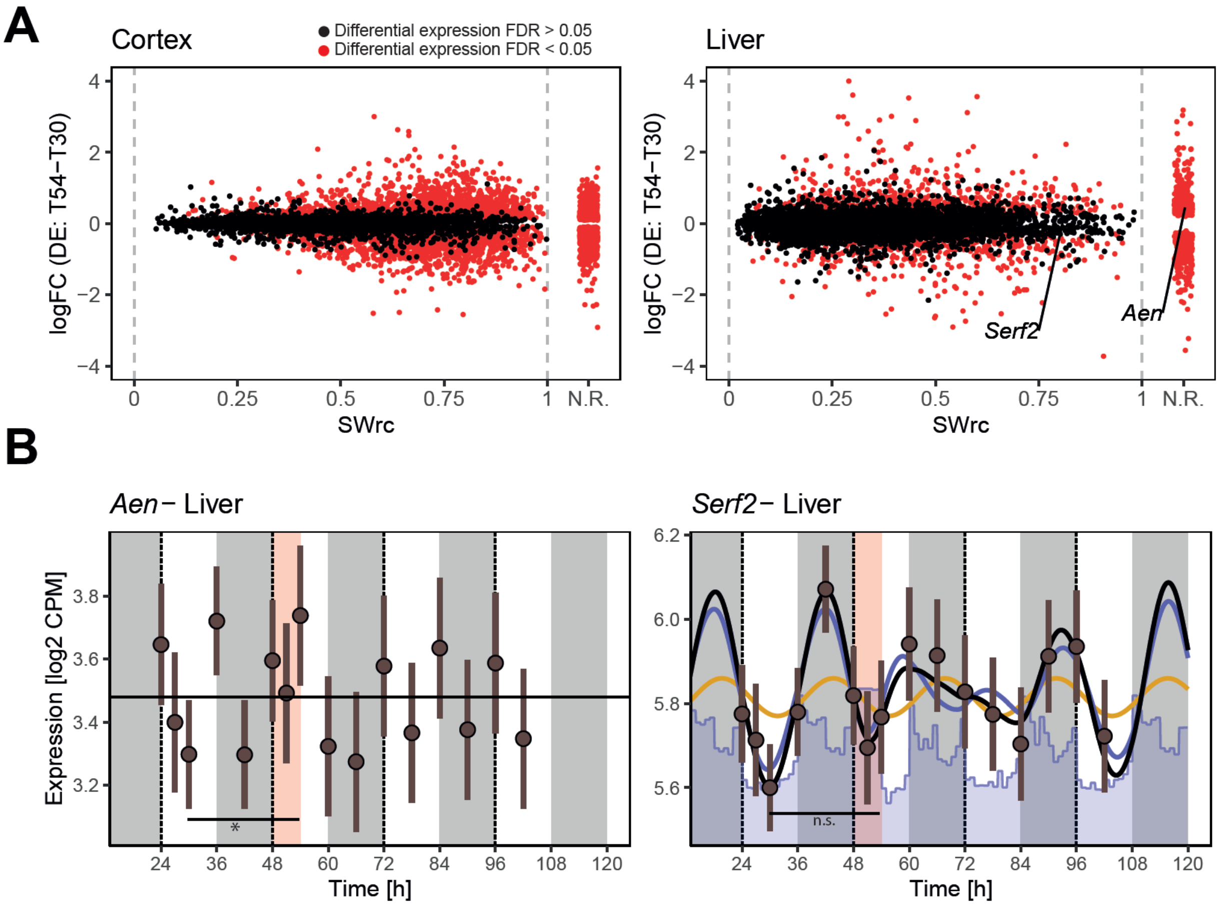


**Figure S2: Differential expression versus dynamic modeling. (A)** Union of gene sets found significantly differentially expressed (DE, red dots) at the end of sleep-deprivation (SD $T54_{ZT6}$ vs. $T30_{ZT6}$) and genes found rhythmic (**Fig. 4B**, $\Delta BIC_{10} > 2$; i.e., $\mathrm{DE} \cup \mathrm{Rhythmic}$), in cortex (left) and liver (right panel). On x-axis the model derived sleep-wake response contribution (*SWrc*) for all rhythmic genes. Non-rhythmic genes (N.R., $\Delta BIC_{10} < 2$) found to be DE ($\mathrm{DE} \cap \mathrm{Rhythmic}'$). Black dots are rhythmic genes found not to be DE ($\mathrm{DE}' \cap \mathrm{Rhythmic}$). **(B)** *Aen* liver expression as example of ($\mathrm{DE} \cap \mathrm{Rhythmic}'$) and *Serf2* illustrating ($\mathrm{DE}' \cap \mathrm{Rhythmic}$), with sleep-wake response in blue and circadian response yellow.
