## Supplementary material for "Model integration of circadian and sleep-wake driven contributions to rhythmic gene expression reveals novel regulatory principles": Fig. S3

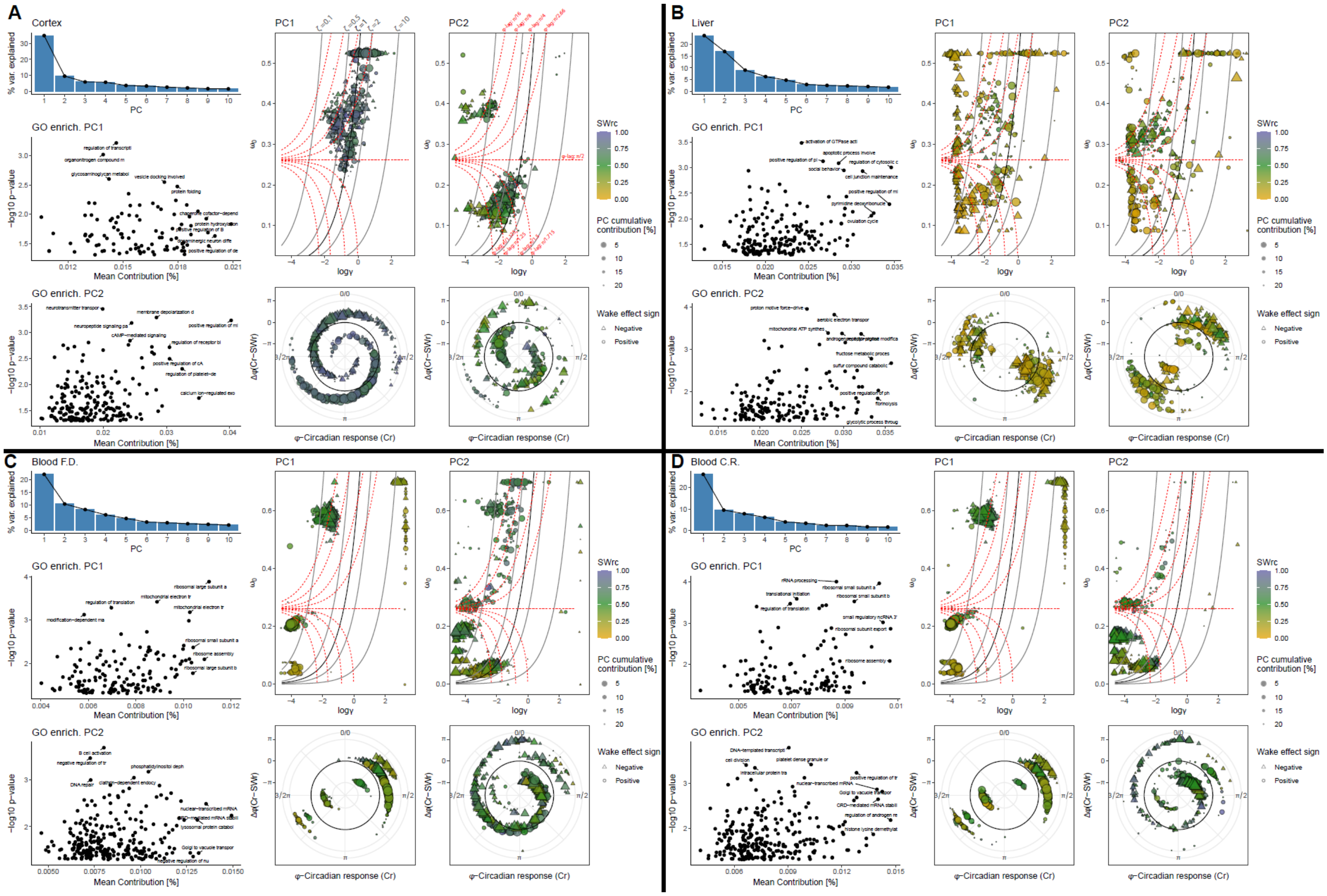

**Figure S3: Supplementary information for the principal component analyses (PCA). (A)** Model parameters and GO term enrichment for PC1 and PC2 in the cortex. Screeplot of the eigenvalues of PCs in the PCA (top-left). GO term enrichment for genes contributing to PC1 (middle-left) and PC2 (bottom-left), x-axis represents mean PCA contribution of genes in the GO-term. Model parameters of genes contributing to PC1 and PC2 up to 20%, with damping (γ) and natural frequency ($\omega_0$) parameters (top-middle, top-right). Black lines represent levels of damping ratio (ζ; units on the top of the PC1 graph). Red lines represent phase-lag between force and response (ϕ-lag; units on the top of the PC2 graph). Phase of the circadian response (Cr) of the model versus the difference between Cr's phase and that of the sleep-wake response (SWr) in baseline (Δϕ[Cr–SWr] for PC1 and PC2 genes, respectively (bottom-middle and -right panels). Black circle represents SWr and Cr being in phase (Δϕ[Cr–SWr] = 0). Symbols represent each gene contributing to PC1 and PC2 with its color indicating the sleep-wake response contribution (*SWrc* range 0-1: yellow-to-blue hues), its size its contribution to the PC (up to 20% of PC cumulated contribution) and its shape indicating whether waking increasing (circle) or decreased (triangle) gene expression rate. Same analyses for data in mouse liver **(B)** and in human blood for the FD **(C)** and CR **(D)** protocols.
