## Supplementary material for "Model integration of circadian and sleep-wake driven contributions to rhythmic gene expression reveals novel regulatory principles": Fig. S4

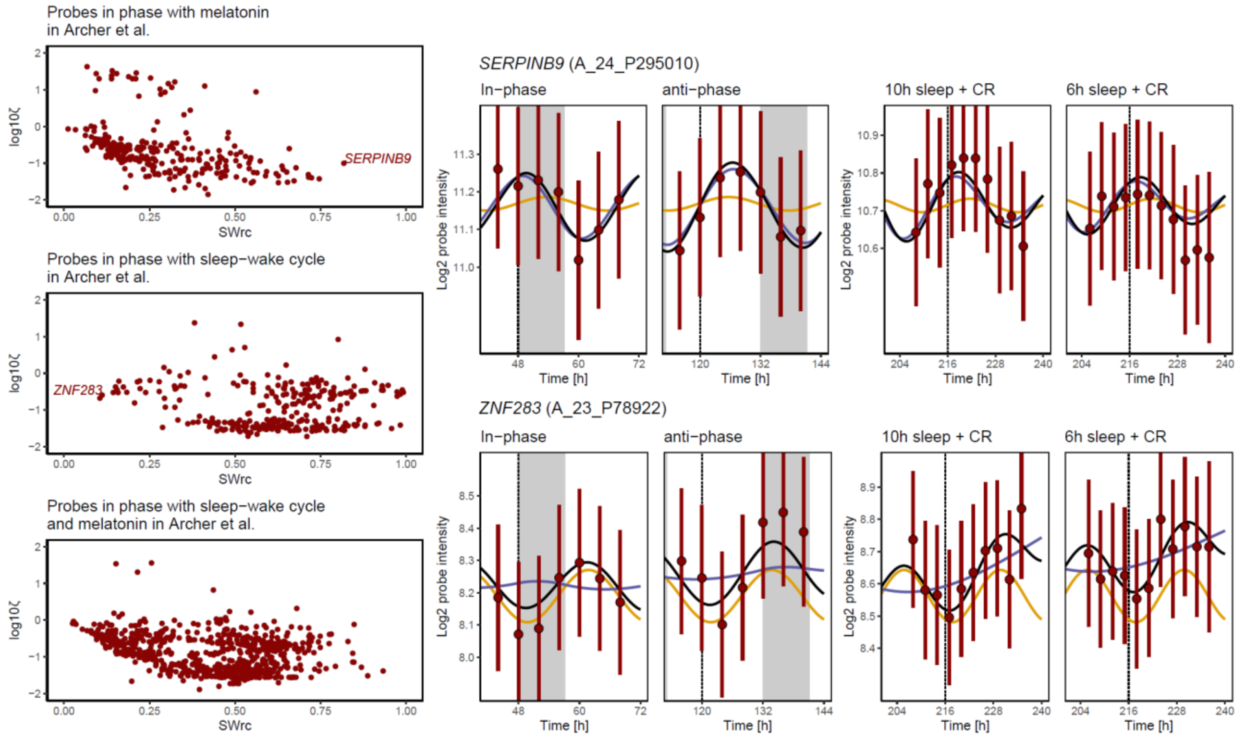


**Figure S4: Compare categorization of genes in Archer *et al.* with our model:** Sleep-wake response contribution (*SWrc*) and damping ratio (ζ) of genes categorized previously in the forced-desynchrony as in-phase with melatonin (top-left), in-phase (middle-left) with sleep-wake cycle, or both (bottom-left panel; (Archer, Laing et al. 2014)). Right panels: Example of a gene categorized as in phase with the circadian melatonin production, that the model, however, identified as a strongly sleep-wake driven (*SERPINB9*; *SWrc* = 0.82; top panels) and of gene categorized as in-phase with the sleep-wake cycle, which was found to be strongly circadian driven (*ZNF283*; *SWrc* = 0.10; lower panels). Vertical red bars span 95% CI of gene expression, blue lines depict the sleep-wake response, yellow lines the circadian response, and black lines represents modelled gene expression.
